# MSTO1 functions as a TRiC assembly factor linking cytosolic proteostasis to mitochondrial function

**DOI:** 10.64898/2026.07.21.739937

**Authors:** Amy M. Bounds, Chloe Higuchi, Suzanne Hoppins

## Abstract

Bi-allelic mutations in *MSTO1* are linked to clinical disease phenotypes characteristic of mitochondrial dysfunction, including ataxia and muscular dystrophy. Consistent with this, MSTO1 patient-derived fibroblasts have fragmented mitochondria and a striking loss of mtDNA. Although MSTO1 has been implicated in regulating mitochondrial fusion, the molecular function of this cytosolic protein in vertebrate cells remains unclear. Using the auxin-inducible degradation (AID) system we demonstrate that MSTO1-FLAG-AID protein is rapidly depleted to almost undetectable levels. Importantly, these cells recapitulate the fragmented mitochondrial phenotype observed in patients and thus are a valuable model of disease. Surprisingly, prior to any changes in mitochondria, we show that MSTO1-depleted cells have a significant decrease in TRiC levels, an essential cytosolic ATP-dependent chaperone required to fold diverse substrates, including actin and tubulin. We reveal that TRiC is also reduced in MSTO1 patient-derived fibroblasts, indicating that loss of TRiC may contribute to disease pathophysiology. We further demonstrate that knockdown of TRiC leads to a decrease in MSTO1 protein levels and remarkably, was sufficient to induce a fragmented mitochondrial phenotype, independent of changes in tubulin or actin. This reveals a previously unrecognized connection between TRiC and mitochondrial homeostasis. Using co-immunoprecipitation we found that MSTO1 interacts with the TRiC chaperone. We also observe accumulation of early TRiC assembly subcomplexes in the absence of MSTO1 suggesting that MSTO1 facilitates assembly of TRiC. Together, our findings identify MSTO1 as a TRiC assembly factor and connect mitochondrial defects caused by MSTO1-depletion to the loss of TRiC.

**Significance:** MSTO1, a protein linked to myopathy and ataxia, has been thought to control mitochondrial fusion, although the molecular mechanism is unknown. Using rapid depletion of MSTO1, we found that mitochondrial fragmentation appears only after six days. Significantly, the levels of the essential cytosolic chaperonin TRiC are reduced within two days of MSTO1 depletion. Directly depleting TRiC reproduces the fragmented mitochondrial phenotype seen with loss of MSTO1, consistent with a model where mitochondrial dysfunction is a downstream consequence of impaired protein folding rather than a direct effect of MSTO1 loss. We show that MSTO1 is required for assembly of TRiC, identifying it as a long-sought assembly factor for this macromolecular protein complex.

## Introduction

Mitochondria contribute to a plethora of cellular processes including energy production, lipid and amino acid biosynthesis, innate immune responses and commitment to cell death. To fulfill these roles, mitochondrial function is not autonomous but relies on coordination with other organelles and cytosolic activities. A defining feature of mitochondria is the maintenance of an ancient and essential genome that encodes components of the oxidative phosphorylation machinery. Mitochondrial dysfunction, resulting from direct or indirect disruption is common in muscular myopathies, heart disease, and neurodegenerative diseases (1–3).

MSTO1 is a cytosolic protein required to maintain mitochondrial structure and function, although molecular details are lacking. Patients with a neuromuscular disease have been documented with compound heterozygous mutations in the nuclear gene, *MSTO1* (4–14). MSTO1 patients present with myopathy-dystrophy, ataxia, cerebellar atrophy, and sometimes retinopathy and cognitive impairments. MSTO1 patient-derived fibroblasts have almost undetectable MSTO1 protein levels, suggesting that reported amino acid substitutions or deletions reduce protein stability (5, 6, 9). Characterization of these cells also implicates MSTO1 in maintaining mitochondrial functions as the cells have a less connected mitochondrial network and fewer copies of the mitochondrial genome (mtDNA). Mitochondrial fusion and fission are critical for maintenance of mitochondrial function and patient-derived fibroblasts have reduced mitochondrial fusion activity compared to control fibroblasts (4, 6). A striking 30-60% decrease in mtDNA content has been reported in several patient-derived fibroblasts (4–6, 8–12). Following MSTO1 knockdown in HeLa cells, mitochondrial motility, membrane potential, and calcium uptake were unaffected, indicating that these mitochondrial defects are not secondary to bioenergetic or movement impairments (4). The molecular functions of MSTO1 remain unknown, and it is unclear how the loss of MSTO1 leads to defects in mitochondrial fusion and mtDNA maintenance.

Interestingly, the primary cellular role of the *Drosophila* MSTO1 homolog Misato is to stabilize the tubulin chaperone protein-1 complex (TCP-1 and TCP-1 ring complex [TRiC in vertebrates]), a cytosolic chaperonin predicted to fold ∼10% of cytosolic proteins, including obligate substrates like tubulin and actin (15, 16). Conditional knockout of *misato* in *Drosophila* larvae reduced the amount of TCP-1 complex, resulting in insufficient amounts of *α*-tubulin to support spindle assembly, ultimately causing mitotic arrest (16). Due to this indirect role in supporting cell division, *misato* is essential for development, and mutant larvae are almost devoid of imaginal disk tissue, have a reduction in brain size, and die before the late third-instar larval stage (15). Also consistent with a role for Misato in TCP-1 stability, tissue specific knockdown of Misato in the intestines resulted in visceral myopathy due to aberrant tubulin and actomyocin structures (17). In conditional brain-specific *misato* knockouts, the mitochondrial network was reported to be similar to controls while fragmentation was reported in the study of visceral muscles. These data imply that loss of *Misato* and TCP-1 in *Drosophila* has tissue-specific effects and that mitochondria can be affected. Of note, both actin and tubulin are required for mitochondrial transport, fission, and fusion and nascent actin and tubulin both require TCP-1/TRiC to reach a final functional folded state (18–23). As demonstrated with the mitotic spindle in *Drosophila*, it is possible that Misato-dependent depletion of TCP-1 in visceral muscle first alters cytoskeletal structures, which then alters mitochondrial structure and/or function. Given that other TCP-1 substrates are involved in a wide range of processes, including cell cycle, translation, transcription, and autophagy, these could also impact mitochondrial function in the absence of Misato (24–30). The possible connection between TCP-1/TRiC and MSTO1 has not been systematically explored in vertebrates, leaving it unknown whether this function of MSTO1 is conserved.

TRiC is an essential 1 MDa hetero-oligomeric ATP-dependent chaperone composed of two rings with 8 subunits each (CCT1-CCT8)(21, 31–33). The precise arrangement of these subunits creates an asymmetric chamber with partitioned biochemical properties required for substrate folding. The mechanism of TRiC assembly remains unresolved. The relative arrangement of the eight paralogous subunits in TRiC are conserved across species and is critical for TRiC function (34–36). Although subunits have only ∼30% sequence identity, the structural similarity is very high and thus presents a challenge to incorporate each subunit in a defined order. Assembly is also critical for stability of the subunits and complex. Excess subunits are degraded, and depletion/loss of any one subunit compromises the stability of the entire complex, resulting in its depletion (35, 37). While each CCT subunit can be purified and is functional based on ATPase activity, they do not assemble into a functional TRiC complex. Rather, some subunits form homooligomers and others interact promiscuously with non-cognate neighbors (38–40). A recent paper outlined a hierarchical pathway for TRiC assembly in mammalian cells where assembly occurs through intermediate scaffolds, and these are rapidly degraded when not in use (35). Calculating predicted free energy of formation ΔG for each subunit interface indicates that the assembly pathway is not spontaneous, suggesting an unknown external regulation drives proper assembly in mammalian cells (35). Together this suggests that efficient TRiC assembly requires chaperones similar to assembly of comparable macromolecular machines like the proteasome or ribosome (41–46). Candidate assembly chaperones have not been identified.

In this study, we find that MSTO1 and TRiC are functionally connected and both are linked to mitochondrial homeostasis. Using an auxin-degradation system in HCT116 cells, we find that MSTO1-depleted cells have fragmented mitochondria, like in MSTO1 patient cells. In this system, we also demonstrate that TRiC protein steady-state levels are decreased days before changes in mitochondrial morphology. Significantly, the same reduction in TRiC is observed in MSTO1 patient-derived fibroblasts, indicating that our system is suitable to inform our understanding of disease pathophysiology. We establish TRiC is required to maintain MSTO1 protein levels and that knockdown of TRiC is sufficient to alter mitochondrial morphology, consistent with a functional connection between TRiC and mitochondria that may be mediated by MSTO1. Consistent with this, we demonstrate that MSTO1 interacts with TRiC and loss of MSTO1 leads to accumulation of TRiC early assembly subcomplexes, implicating MSTO1 in early steps of TRiC assembly. Overall, our data suggest MSTO1 is a novel TRiC assembly factor that acts as a chaperone for subunits to promote assembly into the complex.

## Results

### MSTO1-depleted cells have fragmented mitochondria

To capture the cellular consequences of depletion of MSTO1 with temporal information, we built HCTT16 cells expressing an auxin-inducible degradation (AID) system to control the expression of MSTO1 (47). First, OsTIR1-F74G-V5, the ubiquitin ligase required to identify the AID degron (hereafter OsTIR1-V5), was integrated into HCT116 cells (48, 49). Subsequently, the C-terminus of the endogenous *MSTO1* locus was engineered to encode a FLAG-tag, for detection and downstream biochemical experiments, followed by the mAID-tag. Expression of MSTO1-FLAG-AID was confirmed by Western blot analysis, and knock-in at both alleles was confirmed by PCR (Figure 1A and S1). Of note, recent duplication of a discrete region on chromosome 1q22 in human cells created *MSTO2P*, a pseudogene of *MSTO1* that produces a long noncoding RNA (lncRNA)(50). Due to the very high sequence identity between *MSTO1* and *MSTO2P,* integration of the FLAG-AID sequence also occurred at the 3-prime end of *MSTO2P*. *MSTO2P* is not predicted to make functional protein as a frame shift mutation that leads to a premature stop codon is found in *MSTO2P* before the last exon; therefore, the FLAG-AID insertion in *MSTO2P* is not expected to affect our results.

**Figure 1.**
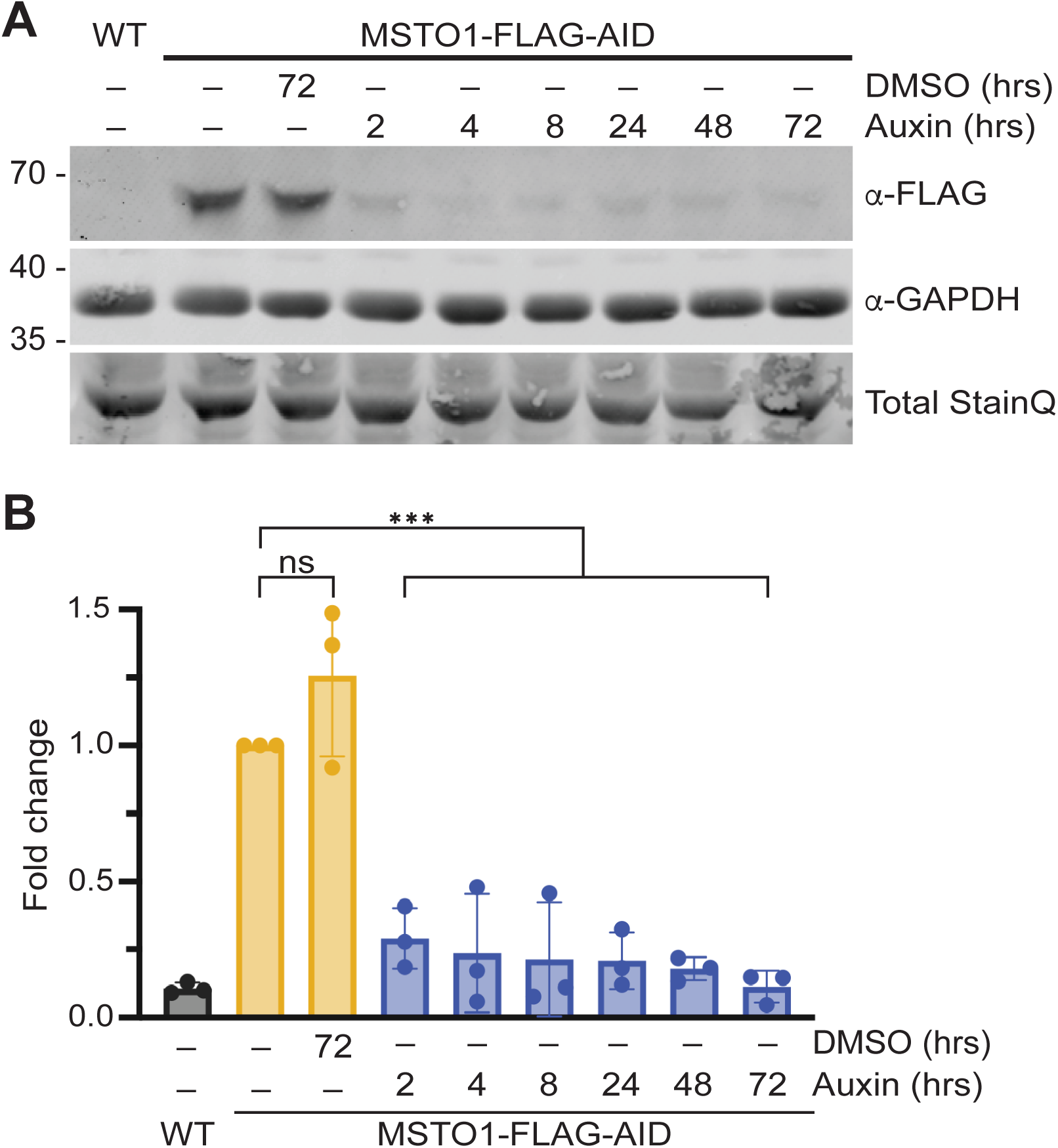
MSTO1 is depleted after 2 hours of auxin treatment. **(A)** Representative Western blot showing depletion of MSTO1-FLAG-AID. Whole cell lysate from wild-type HCT116 (WT) and MSTO1-FLAG-AID cells treated with either vehicle (DMSO) or 20 *μ*M auxin for the indicated time were subject to SDS-PAGE and immunoblotting with anti-FLAG and anti-GADPH with Total StainQ used as loading control. Molecular weight markers are indicated in kDa on the left. **(B)** Quantification of Western blot as shown in (A). Fold change of MSTO1-FLAG-AID protein level relative to untreated MSTO1-FLAG-AID cells was calculated. The graph shows the mean and standard deviation from three independent experiments, ns: not significant, ***P < 0.001, ordinary one-way Anova with multiple comparisons.

MSTO1-FLAG-AID was rapidly degraded after addition of the auxin analog, 5-Ph-IAA (hereafter auxin)(Figure 1). Immunoblotting for FLAG revealed that the MSTO1 protein level is 25% of vehicle (DMSO) treated controls after two hours of auxin treatment (Figure 1). This was consistent across three different clonal populations of cells with MSTO1-FLAG-AID (Figure S2). Our MSTO1-FLAG-AID cells offer an unprecedented opportunity to establish the consequences of MSTO1 depletion with temporal resolution unavailable with other approaches to gain insight into the molecular functions of this understudied protein.

We first examined mitochondrial morphology following depletion of MSTO1. We expected to observe an increase in short and/or fragmented mitochondria, as reported in patient-derived fibroblasts (4–12). Mitochondrial morphology was scored in blinded experiments. The mitochondrial network of wild-type HCT116 cells is a mixture of reticular and short, with most mitochondria having a length that is greater than 5 μm, and few mitochondria in the network are between 2.5 and 5 μm in length, which we classify as fragmented. Quantification of mitochondrial morphology in HCT116 cells expressing OsTIR1-V5 yielded results comparable to unmodified wild-type cells. We treated MSTO1-FLAG-AID cells with either vehicle or 20 micromolar auxin for either 24 or 72 hours before imaging. No changes in mitochondrial morphology were observed at these timepoints (data not shown). We extended the time with auxin and found that after six days, MSTO1-depleted cells had a higher percentage of fragmented mitochondria compared to earlier timepoints and vehicle controls (Figure 2A-B). This phenotype was also consistent across three MSTO1-FLAG-AID clonal populations (Figure S3). Previously MSTO1 was knocked down in HeLa cells, where changes in mitochondrial morphology were observed 72 hours after transduction (4, 51). Of note, HeLa cells have multiple copies of chromosome 1 resulting in relatively high expression levels of MSTO1 protein. This unique context may underlie the differences in our data.

**Figure 2.**
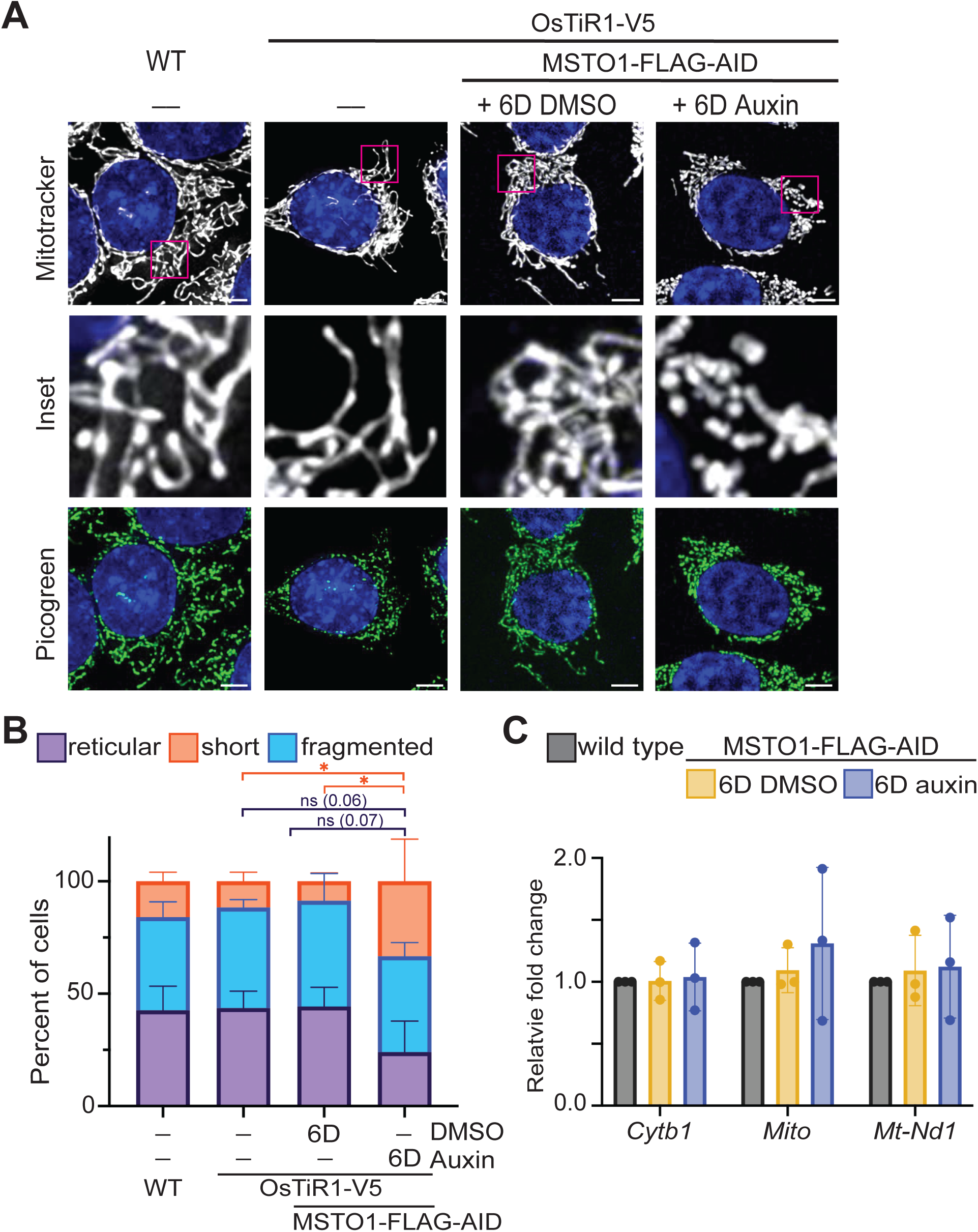
MSTO1-depleted cells have fragmented mitochondria. **(A)** Representative live cell images of wild-type HCT116 (WT), WT+OsTiR1-V5, and MSTO1-FLAG-AID treated with DMSO (vehicle) or 20 *μ*M auxin for 6 days. Mitochondria were labeled with Mitotracker Red CMXRos, mtDNA were labeled with Quant-iT™ PicoGreen® dsDNA Reagent, and nuclei were labeled with NucBlue. All were visualized by fluorescence microscopy. Scale bar is 5 *μ*M **(B)** Quantification of mitochondrial morphology in cell lines described in (A). The graph shows mean of at least 100 cells and standard deviation from three independent experiments. ns: not significant, *P < 0.05, ordinary two-way Anova with multiple comparisons. **(C)** Quantification of mtDNA in MSTO1-FLAG-AID cells treated with vehicle or 20 *μ*M auxin for 6 days. Cellular DNA samples were prepared from the indicated cell line and mtDNA was quantified by qPCR relative to *GAPDH*, a nuclear housekeeping gene. The graph shows the mean and standard deviation of three independent experiments.

Another common observation in MSTO1 patient-derived fibroblasts was a decrease in mtDNA copy number compared to controls. We assessed mtDNA content in our cells in two ways: first by visualizing nucleoids with PicoGreen staining and second with qPCR. We assessed mtDNA following 6 days of auxin treatment, when we observed changes in mitochondrial connectivity. The MSTO1-FLAG-AID cells stained with Picogreen had no obvious changes in mtDNA nucleoid number, size, or distribution compared to wild-type or MSTO1-FLAG-AID cells treated with vehicle (Figure 2A). The relative mtDNA copy number quantified by qPCR was also comparable to controls (Figure 2C). Therefore, in this timeframe, we do not observe a decrease in mtDNA following depletion of MSTO1.

### TRiC levels are significantly reduced before mitochondrial morphology changes

One important step forward in understanding the cellular roles of MSTO1 is to determine if vertebrate MSTO1 is also functionally connected to TRiC, as reported for Misato and TCP-1 (15, 16). Loss of any individual TRiC subunit results in instability of the entire complex and degradation of remaining subunits by the proteasome or autophagy (16, 35). Hence, we use CCT1, the most frequently used antibody to detect the TRiC complex and to monitor TRiC levels following depletion of MSTO1-FLAG-AID. Using the same approach as described in Figure 1, we probed the indicated samples with anti-CCT1 and observed that after 48 hours of auxin treatment, CCT1 protein level was significantly decreased compared to vehicle controls (Figure 3A-B). Similar reduction of CCT1 was observed in all three MSTO1-FLAG-AID isolates (Figure S4A-D). Given that mitochondrial morphology was not altered until six days following depletion of MSTO1, we also quantified CCT1 protein abundance at this timepoint. Indeed, CCT1 protein levels were reduced to ∼40% of control levels after both three and six days of treatment (Figure S4E-F). These data are consistent with previously published data indicating Misato was required for TCP-1 stability and establish the conservation of this function for MSTO1.

**Figure 3.**
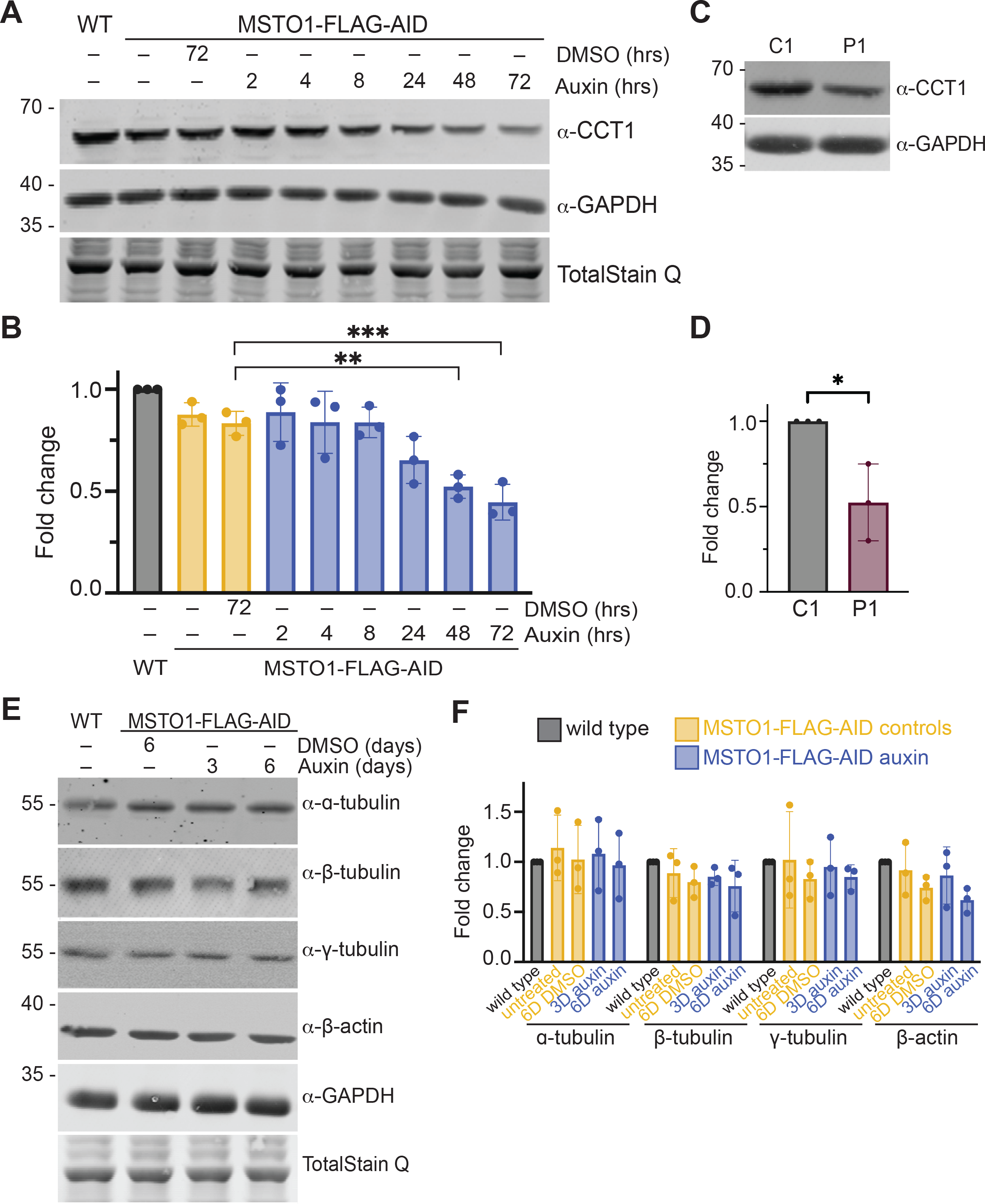
TRiC levels are significantly reduced before mitochondrial morphology changes. **(A)** Representative Western blot showing protein levels of CCT1 following depletion of MSTO1-FLAG-AID. Whole cell lysates from wild-type HCT116 (WT) or MSTO1-FLAG-AID cells treated with either vehicle (DMSO) or 20 *μ*M auxin for the indicated time were subject to SDS-PAGE and immunoblotting with anti-CCT1, and anti-GADPH with Total StainQ used as loading control. Molecular weight markers are indicated in kDa on the left. **(B)** Quantification of Western blot as shown in (A). Fold change of CCT1 protein level relative to WT cells was calculated. The graph shows the mean and standard deviation from 3 independent experiments, **P < 0.01, ***P < 0.001, ordinary one-way Anova with multiple comparisons. **(C)** Representative Western blot showing CCT1 protein levels in patient cells compared to control. Whole-cell lysates prepared from either control or MSTO1 patient fibroblasts were subject to SDS-PAGE and immunoblotting with anti-CCT1 and anti-GADPH. Molecular weight markers are indicated in kDa on the left. **(D)** Quantification of Western blot as shown in (C). Fold change of CCT1 protein levels relative to control patient fibroblasts was calculated. The graph shows the mean and standard deviation from 3 independent experiments, *P < 0.05, Welch’s unpaired T-test. **(E)** Representative Western blot showing protein levels of cytoskeleton components. Whole cell lysates from wild-type HCT116 (WT) or MSTO1-FLAG-AID cells treated with either vehicle (DMSO) or 20 *μ*M auxin for the indicated time were subject to SDS–PAGE and immunoblotting with anti-*α*-tubulin, anti-*β*-tubulin, anti-*γ*-tubulin, anti-*β*-actin, and anti-GADPH with Total StainQ used as loading control. Molecular weight markers are indicated in kDa on the left. **(F)** Quantification of Western blot as shown in (E). Fold change of *α* -tubulin, *β*-tubulin, *γ*-tubulin, *β*-actin protein levels relative to WT HCT116 cells was calculated. The graph shows the mean and standard deviation from 3 independent experiments.

TRiC expression has never been assessed in MSTO1 patients, so we wanted to determine if the loss of TRiC observed in our MSTO1-FLAG-AID system was also a feature of MSTO1 patient-derived fibroblasts. We obtained MSTO1 patient fibroblasts, which have a drastic reduction in MSTO1 protein levels (6). As above, CCT1 steady-state protein levels were quantified as a proxy for the TRiC complex. Our data show that the MSTO1 patient-derived cells have a significant decrease in CCT1 protein level compared to control fibroblasts (Figure 3C-D). This decrease is similar to what we observed in MSTO1-depleted cells, suggesting the MSTO1-FLAG-AID cells are a valid model to understand the relationship between MSTO1, TRiC, and mitochondrial homeostasis.

TRiC is most well-known for its obligate role in folding nascent tubulin and actin (52, 53). Both cytoskeletal elements contribute to mitochondrial morphology and the dynamic processes of transport, fusion, and fission (20). Long-range mitochondrial movements are microtubule-based, mediated by dynein and kinesin (54, 55). Actin has diverse roles in regulating mitochondrial dynamics, both through myosin-mediated movement and actin polymerization (56–60). Actin also plays an active role in mitochondrial fission by promoting constriction and recruiting Drp1 (19, 61, 62). Actin has also been visualized at mitochondrial fusion sites, but its mechanistic role here remains unclear (23). Therefore, we hypothesized that if tubulin and/or actin steady-state protein levels were decreased due to the loss of TRiC, this could impact mitochondrial connectivity.

To test this, we quantified *α*-, *β*-, *γ*-tubulin, and *β*-actin protein steady-state levels after three and six days of auxin treatment to capture the consequence of the depletion of MSTO1 and TRiC, which are decreased at two hours and two days, respectively. Despite the significant decrease of TRiC, no changes in steady-state levels of any of the cytoskeletal components was observed at either timepoint (Figure 3E-F). This was also consistent across two MSTO1-AID-FLAG isolates (Figure S5A-B). Accordant with this, immunofluorescence analyses following depletion of MSTO1 and TRiC revealed no clear changes in the microtubule or actin cellular networks (Figure S5C). These data suggest that there is sufficient TRiC in MSTO1-depleted HCT116 cells to maintain steady state levels of tubulin and actin and their respective cytoskeletal structures.

### TRiC is required to maintain MSTO1 steady state protein levels and mitochondrial morphology

To date, there is no evidence in vertebrates directly linking TRiC chaperone function with mitochondrial homeostasis, but our data suggest that TRiC may be required to maintain mitochondrial structure. To test this, we reduced CCT1 protein levels by expression of short hairpin RNA (shRNA) targeting CCT1 transcripts in MSTO1-FLAG-AID cells. As discussed earlier, loss of CCT1 has been shown to reduce the amount of assembled TRiC; therefore, this condition will deplete the TRiC complex. Compared to the control shRNA targeting RFP, cells expressing shCCT1 A or shCCT1 C had reduced CCT1 protein levels (∼40% and ∼60% respectively)(Figure 4A-B). To determine the impact of decreased TRiC on mitochondrial function, we first scored mitochondrial morphology in cells expressing either of our CCT1-targeting shRNAs. In both cases, we quantified an increased portion of the population with fragmented and short mitochondria compared to controls (Figure 4C-D). Next, we examined mitochondrial nucleoids, which revealed no changes in mtDNA nucleoid number or distribution (Figure 4C). Therefore, loss of TRiC leads to altered mitochondrial morphology and the change is strikingly similar to what has been observed in MSTO1 patients and our MSTO1-depleted cells.

**Figure 4.**
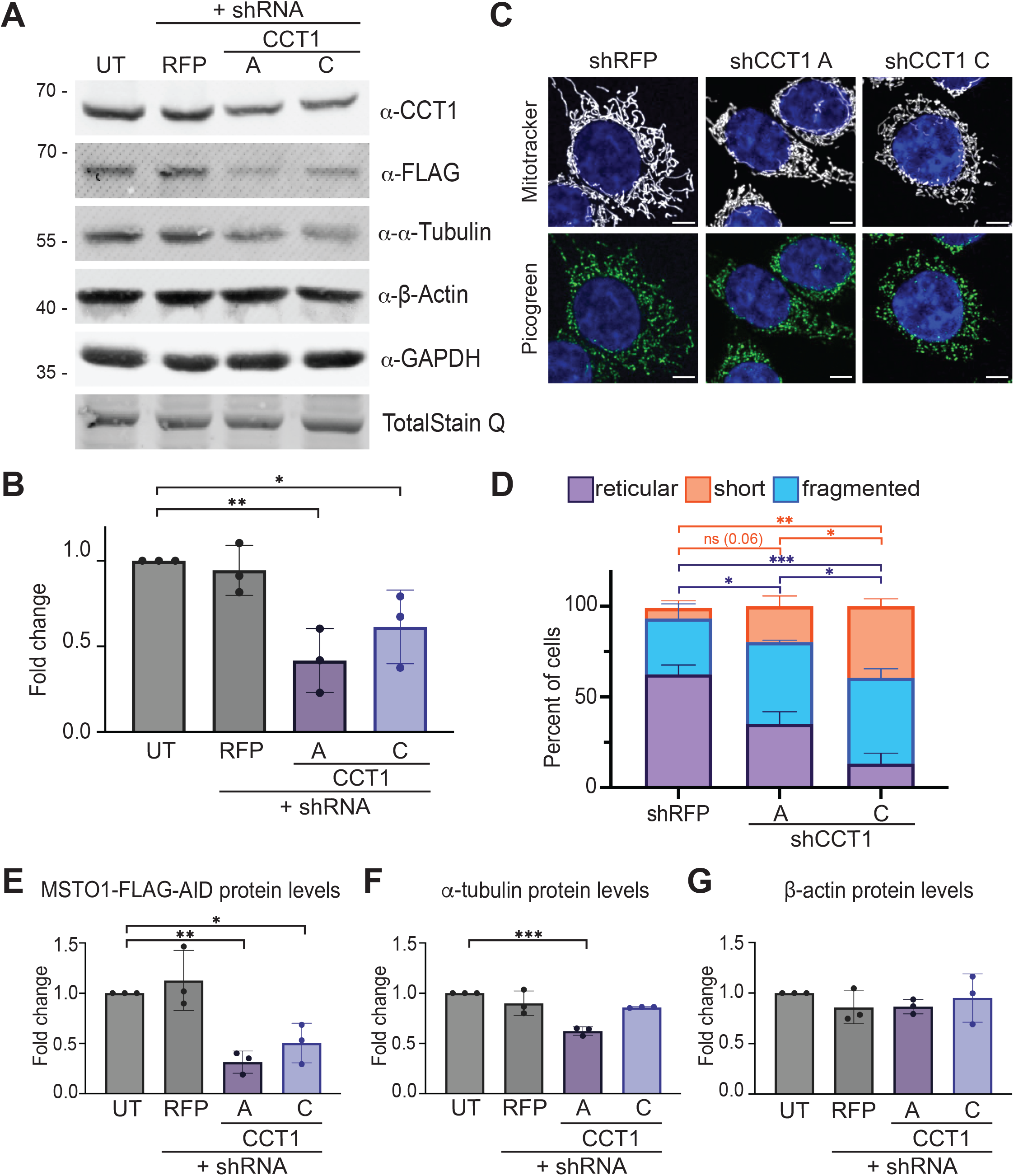
TRiC is required to maintain MSTO1 steady state protein levels and mitochondrial morphology. MSTO1-FLAG-AID cells expressing either control shRNA (shRFP) or shRNAs targeting MSTO1 (shCCT1 A or C). **(A)** Representative Western blot showing protein levels following knockdown of CCT1. Whole-cell lysates prepared from MSTO1-FLAG-AID cells that were untreated (UT) or expressing shRFP, shCCT1 A, or shCCT1 C were subject to SDS-PAGE and immunoblotting with anti-CCT1, anti-FLAG, anti-*α*-tubulin, anti-*β*-actin, and anti-GADPH with Total StainQ used as loading control. Molecular weight markers are indicated in kDa on the left. **(B)** Quantification of Western blot as shown in (A). Fold change of CCT1 protein level relative to untreated cells was calculated. The graph shows the mean and standard deviation from 3 independent experiments, *P < 0.05, **P < 0.01, ordinary one-way Anova with multiple comparisons. **(C)** Representative live cell images of MSTO1-FLAG-AID +shRFP, +shCCT1 A, or +CCT1 C. Mitochondria were labeled with Mitotracker Red CMXRos, mtDNA were labeled with Quant-iT™ PicoGreen® dsDNA Reagent, and nuclei were labeled with NucBlue. All were visualized by fluorescence microscopy. Scale bar is 5 *μ*M. **(D)** Quantification of mitochondrial morphology for cell lines described in (C). The graph shows the mean and standard deviation from 3 independent experiments *P<0.05, **P < 0.01, ***P < 0.001, ordinary two-way Anova with multiple comparisons. **(E-G)** Quantification of Western blot as shown in (A). Fold change of FLAG (E), *α*-tubulin (F), or *β*-actin (G) protein levels were calculated. The graph shows mean and standard deviation from 3 independent experiments *P<0.05, **P < 0.01, ***P < 0.001, ordinary one-way Anova with multiple comparisons.

As shown above, MSTO1 is required to maintain TRiC protein levels. To determine if TRiC in turn regulates the steady state levels of MSTO1, we assessed MSTO1-FLAG protein levels following CCT1 knockdown. In both knockdowns of CCT1, we observe significant decreases in MSTO1-FLAG-AID protein (Figure 4A and E, 30% and 43% of controls, respectively). Therefore, in humans, MSTO1 stability depends on TRiC and vice versa. It is important to note that since MSTO1 protein levels are reduced in TRiC-depleted cells, it is impossible to deconvolve whether the change in mitochondrial morphology is due to the loss of MSTO1, TRiC, or both.

We next sought to determine if steady state levels of either tubulin or actin are affected by the reduced levels of TRiC in our CCT1-knockdown conditions. We quantified *α*-tubulin and *β*-actin by Western blot in cells expressing shRNAs targeting RFP or CCT1 (Figure 4A and F-G). We find that steady state *β*-actin levels are unchanged following depletion of TRiC by either shRNA targeting CCT1 (Figure 4A and F-G). In one of the knockdown conditions, *α*-tubulin levels were decreased compared to controls (Figure 4A and F, shRNA A). The difference between the two shRNA lines is consistent with the greater reduction of CCT1 levels with the shCCT1 C shRNA. Together, our data demonstrate that shRNA-directed depletion of TRiC phenocopies AID-dependent depletion of MSTO1 and the impact on mitochondrial morphology.

### MSTO1 physically interacts with TRiC

Our data reveal that MSTO1 is required for TRiC stability and *vice versa*. Previously, MSTO1 has been suggested to regulate mitochondrial fusion, but it remains unclear whether MSTO1 directly modulates membrane fusion activity. These functions suggest that MSTO1 may physically interact with the outer mitochondrial membrane fusion machine and/or TRiC. To test this, we immunoprecipitated the endogenously tagged MSTO1-FLAG-AID and used mCherry-FLAG expressing wild-type HCT116 cells as a negative control to demonstrate specific binding (Figure 5A). The immunoprecipitation of MSTO1-FLAG-AID was efficient. While neither Mfn1 nor Mfn2 interacted with MSTO1 in this assay, we found that endogenous MSTO1 does interact with endogenous CCT1.

**Figure 5.**
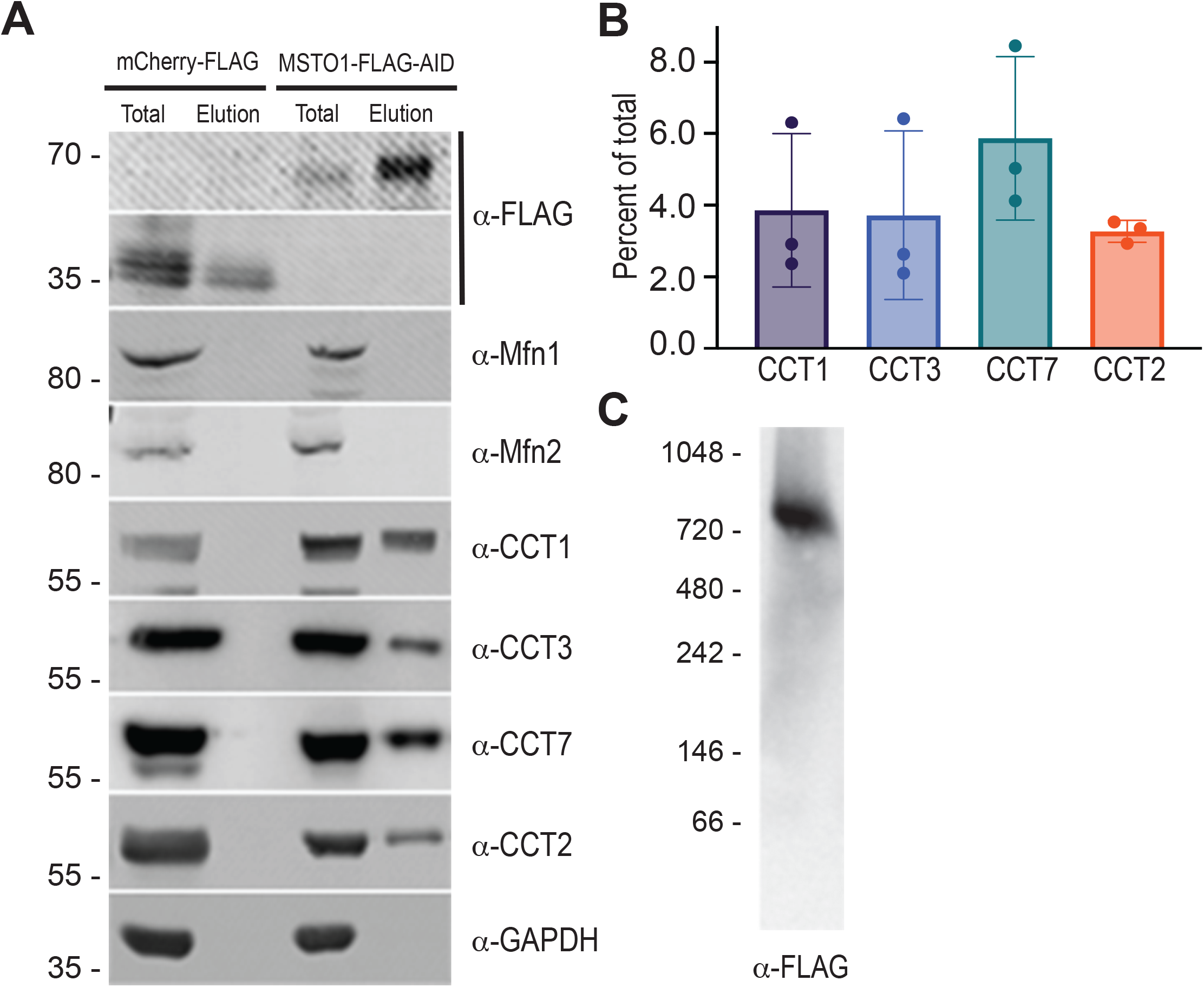
MSTO1 interacts with TRiC. **(A)** Representative Western blot of immunoprecipitation of either control (mCherry-FLAG) or MSTO1-FLAG-AID using anti-FLAG magnetic beads. Lysate fraction (Total, 0.66%) and immunoprecipitates (Elution, 6.6%) were subject to SDS-PAGE and immunoblotting with anti-FLAG, anti-Mfn1, anti-Mfn2, anti-CCT1, anti-CCT3, anti-CCT7, anti-CCT2, or anti-GADPH. Molecular weight markers are indicated in kDa on the left. **(B)** Quantification of co-immunoprecipitated CCT1, CCT3, CCT7, and CCT2 protein levels compared to total protein levels from Western blot in (A). **(C)** Representative blue native polyacrylamide gel electrophoresis (BN-PAGE) of MSTO1-FLAG-AID. Whole cell lysates from MSTO1-FLAG-AID cells treated vehicle for 3 days were subject to BN-PAGE followed by immunoblotting with anti-FLAG. Molecular weight markers are indicated in kDa on the left.

The vast majority of CCT subunits are found in fully assembled TRiC complexes and orphaned subunits are rapidly degraded by quality control pathways (37). However, recent reports indicate that low levels of CCT subcomplexes exist at steady state, specifically CCT2-4, CCT5-7 and CCT1-3-6 (35). Therefore, it is possible that MSTO1 interacts with either the holocomplex or one or more subcomplexes. To test this, we chose one subunit from each early assembly subcomplex and tested for interaction with MSTO1 in our co-immunoprecipitation assay. We find that endogenous MSTO1 interacts with endogenous CCT2, CCT7, and CCT3, representing each early assembly subcomplex (Figure 5A-B). The results were consistent across two clonal populations of MSTO1-FLAG-AID cells (Figure S6A-B). The efficiency of co-immunoprecipitation was similar for all CCT subunits tested. Therefore, this assay does not distinguish between interactions of MSTO1 early assembly subcomplex or fully assembled TRiC.

If MSTO1 interacts with the TRiC complex, we would expect it to exist as a high molecular weight species, as previously reported for TRiC (35). Therefore, we sought to determine the relative size of MSTO1 in cells. Lysates from MSTO1-FLAG-AID cells treated with vehicle were separated by Blue Native-PAGE (BN-PAGE) and immunoblotted with anti-FLAG. Consistent with its association with TRiC, MSTO1-FLAG-AID is only detected in a high molecular weight complex (Figure 5C, Figure S6C). While it is possible that MSTO1 self-assembles into a homo-oligomer of this size, this is also consistent with MSTO1 stably interacting with TRiC, as indicated by our co-immunoprecipitation data where every CCT subunit tested interacted with MSTO1.

Interestingly, MSTO1 and TRiC subunits are estimated to have drastically different cellular concentrations or protein copy number. These relative concentrations were calculated by quantitative mass spectrometry analysis of endogenously tagged genes in HEK293T cells (63). TRiC is known to be abundant, and this report calculated that each TRiC subunit is present at ∼6000 nM in human cells. In contrast, MSTO1 protein level measurements indicate concentrations about 20-fold less abundant (∼170 nM). Thus, it is unlikely that MSTO1 is a stoichiometric interaction partner with all TRiC macromolecular complexes.

### MSTO1 is a novel TRiC assembly chaperone

All factors required for the assembly of TRiC have not been identified. Within the TRiC complex, the CCT subunits are arranged in a precise order that is required for function and stability of the complex. The subunits cannot spontaneously assemble, and unassembled orphan subunits are rapidly degraded (37). Assembled TRiC can be visualized by native gel electrophoresis, which also revealed low levels of early assembly subcomplexes that accumulate in cells where TRiC assembly is compromised (35). The current model for TRiC assembly posits that CCT2 and CCT4 interact to form the earliest assembly subcomplex. To this unit, a subcomplex composed of CCT5 and CCT7 is added followed by CCT8 alone and finally a subcomplex with CCT1, CCT3, and CCT6. To determine if MSTO1 plays a role in TRiC assembly, MSTO1-FLAG-AID cells were treated with auxin for three days and cell lysates were subject to BN-PAGE and immunoblotting against CCT3, CCT7, and CCT2 to capture the assembly subcomplexes that were previously identified (35, 64). In control conditions, we observe both fully assembled TRiC and a small proportion of CCT3, CCT7 and CCT2 in smaller subcomplexes, consistent with previous reports (Figure 6 and S7)(35). In keeping with our data presented in Figure 3, following three days of auxin treatment, fully assembled TRiC was decreased in all cases (Figure 6, closed arrowhead). Interestingly, following the depletion of MSTO1, there is an accumulation of CCT2 and CCT7, which interact with CCT4 and CCT5 respectively in distinct subcomplexes (Figure 6 and S7A-B, open arrowhead). In contrast, there is no change in the subcomplexes containing CCT3, which is predicted to be the last subcomplex to assemble into the ring. These data were consistent across two clonal populations of MSTO1-FLAG-AID (Figure S7). The accumulation of the first two assembly subcomplexes in the absence of MSTO1 is consistent with a model where MSTO1 facilitates assembly of the CCT2-CCT4 and CCT5-CCT7 subcomplexes which would then be used as a scaffold for assembly of CCT8 and the subcomplex composed of CCT1-3-7.

**Figure 6.**
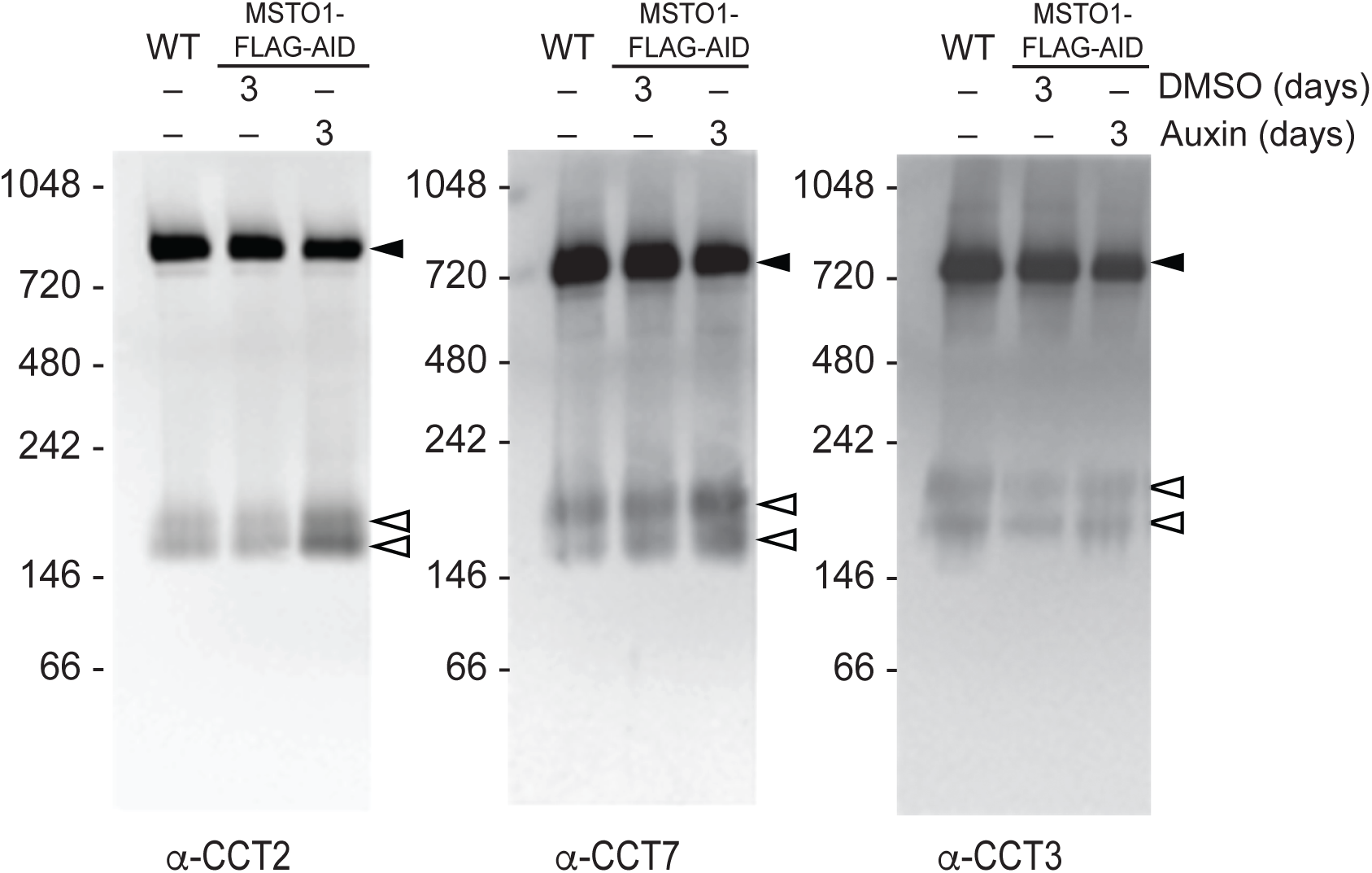
MSTO1 facilitates assembly of TRiC. **(A)** Representative BN-PAGE showing fully assembled TRiC and early assembly scaffolds following depletion of MSTO1-FLAG-AID. Whole cell lysates from untreated wild-type HCT116 (WT) or MSTO1-FLAG-AID cells treated with vehicle (DMSO) or 20 *μ*M auxin for 3 days were subject to BN-PAGE followed by immunoblotting with anti-CCT2, anti-CCT3, or anti-CCT7. Open arrowhead indicates early assembly scaffolds and closed arrowhead indicates fully assemble TRiC. Molecular weight markers are indicated in kDa on the left.

## Discussion

MSTO1 is primarily cited as a cytosolic protein required for mitochondrial fusion. This was based on reports of bi-allelic pathogenic variants of *MSTO1* resulting in congenital or early onset mitochondrial myopathy and ataxia and the study of MSTO1 patient-derived fibroblasts and HeLa cells with reduced MSTO1 expression. These cells had reduced mitochondrial fusion activity and mtDNA copy number. Interestingly, this was not consistent with characterization of the *Drosophila* homolog, Misato, which was required for stability of the cytosolic chaperonin TCP-1/TRiC (16). Here, depletion of Misato resulted in embryonic lethality due to cell cycle defects. Our data puts forth a unifying model of MSTO1 using an AID system to temporally control the expression of MSTO1. We demonstrate that depletion of MSTO1-FLAG-AID recapitulates the fragmented mitochondrial phenotypes reported in MSTO1 patient fibroblasts. However, depletion of MSTO1 in our system did not result in loss of mtDNA, which has been reported in many MSTO1 patient-derived cells. This is consistent with a study where reintroduction of MSTO1 to patient fibroblasts rescued mitochondrial morphology but not mtDNA depletion after 48 hours (9). Together with our data, this suggests that mtDNA loss in patients is a more complex process rather than a direct result of MSTO1 depletion. It may be that mtDNA loss is gradual and not evident after six days of MSTO1 depletion or that the HCT116 cells do not fully recapitulate a physiological or developmental state that strictly requires MSTO1 for mtDNA maintenance.

Our data are not consistent with MSTO1 directly regulating the mitochondrial fusion machinery. Changes in mitochondrial structure were observed only after MSTO1 was depleted for six days. Given that mitochondrial fusion occurs once a minute or more in cell culture conditions (65), this timescale suggests that loss of MSTO1 does not acutely impact mitochondrial fusion rates. Furthermore, MSTO1 does not interact with either mitofusin protein. Together, these data challenge the model that the molecular function of MSTO1 is to directly regulate mitochondrial fusion and prompted our investigation of a potential connection between MSTO1 and TRiC.

TRiC is best known for its obligate role in folding tubulin and actin, although its clients include many essential proteins including F-box proteins, DNA and RNA replication factors, cell cycle proteins, subunits of mTORC, and protein kinases (24–30, 66). Although the MSTO1 and TRiC *Drosophila* homologs have been functionally connected, the role of TRiC in MSTO1-related disease had not been previously assessed. Our experiments reveal for the first time that depletion of vertebrate MSTO1 results in depletion of TRiC before any changes in mitochondrial structure are observed. We further demonstrate that patients with bi-allelic mutations in *MSTO1* have reduced TRiC levels. In our AID system, TRiC levels decrease four days before our cells have fragmented mitochondria, suggesting that changes in mitochondrial morphology are impacted by loss of TRiC. While the most obvious connection between TRiC and mitochondria are actin and tubulin, we find that the steady state level of both are unaltered in MSTO1-depleted cells and the overall cytoskeletal structures are normal.

Our data predicts that depletion of TRiC would phenocopy depletion of MSTO1. Indeed, we demonstrate that cells with reduced TRiC resulting from CCT1-targeted knockdown also have fragmented mitochondria. Thus, loss of TRiC is sufficient to induce mitochondrial morphology defects. An important caveat here is that depletion of TRiC also results in loss of MSTO1; therefore we cannot dissect any distinct functional roles. The molecular basis of the functional connection between MSTO1-TRiC and mitochondrial homeostasis is not yet clear. The simplest model is that TRiC substrates are required to maintain mitochondrial function. TRiC substrates are characterized by aggregation-prone and topologically complex domains and most are small enough to fit inside the folding chamber, although there are recent reports of non-canonical open-state folding substrates that are much larger than tubulin or actin (67, 68). Given that most nuclear-encoded mitochondrial proteins are translocated across mitochondrial membranes in an unfolded state, we find it most likely that the substrate would be cytosolic or associated with the mitochondrial outer membrane.

TRiC is an abundant and large heterooligomeric assembly comprised of eight related but functionally distinct subunits with high structural similarity. This poses a challenge for assembly, which must create two stacked rings with each subunit in a particular arrangement that results in an asymmetrical inner chamber with partitioned biochemical properties essential for function (33, 35, 69, 70). While co-expression of the eight TRiC subunits in insect cells results in assembly of the complex, combining independently expressed subunits ex vivo does not (71, 72). Furthermore, analysis indicates that the assembly pathway is not driven by the hierarchy of binding free energies (35). Therefore, assembly of TRiC is thought to require an elusive cellular component that promotes on-pathway interactions of the subunits. Our data indicate that MSTO1 is a TRiC assembly factor. Specifically, in the absence of MSTO1, TRiC early assembly subcomplexes (CCT2-CCT4 and CCT5-CCT7) accumulate in the cell, consistent with altered assembly kinetics. The same accumulation of subcomplexes has also been reported when TRiC was depleted by knocking down CCT2 or CCT7 (35). Although not shown in Betancourt Moreira *et al*., it is likely that MSTO1 levels are also reduced in these CCT knockdown conditions as we report here. Therefore, we predict that MSTO1 promotes assembly of two subcomplexes—CCT2-CCT4 and CCT5-CCT7—before addition of CCT1-CCT3-CCT6 and CCT8 to create the fully assembled TRiC. We predict that the accumulation of these subcomplexes is limited due to quality control mechanisms that degrade orphan subunits, which is likely responsible for rapid destruction of CCT1, CCT3, CCT6 and CCT8 in the absence of MSTO1-mediated partial ring assemblies (37).

Our co-immunoprecipitation and BN-PAGE data are consistent with a model where MSTO1 interacts with the fully assembled TRiC complex. We speculate that MSTO1 remains bound to the subcomplexes until the nascent TRiC complex is formed, comparable to chaperones required for proteasome assembly (41, 73). MSTO1 could either be on the outside of the complex and dissociate from the assembled double-ring structure, or it could be encased in the chamber and evicted in a manner similar to other substrates. Given that depletion of TRiC results in lower levels of MSTO1, and the structural similarity between MSTO1 and tubulin, MSTO1 may be a TRiC substrate in addition to an assembly chaperone.

While our data indicate that MSTO1 and TRiC are functionally connected, there are clear indications that their functions do not completely overlap. Most strikingly, patients with mutations in MSTO1 are distinct from patients with mutations in CCT subunits, which were recently coined TRiCopathies (74). TRiCopathy patients present with severe global developmental disorders, brain malformations, microcephaly, muscle hypotonia, ataxia, and intellectual disability (74). This clinical divergence indicates that there is something unique about the loss of MSTO1 that results in mitochondrial dysfunction and present as myopathy and ataxia. In addition, we noticed that cells expressing shRNA that targets CCT1 have cell proliferation defects compared to shRFP controls. In contrast, we never observe slow growth of MSTO1-FLAG-AID cells treated with auxin. Together, these findings suggest that MSTO1 is not simply required for TRiC assembly, but also performs functions that more directly impact mitochondrial function.

## Materials and Methods

### Cell Culture

HCT116 cells were cultured in McCoy’s 5A medium supplemented with 10% FBS (Seradigm), 2 mM L-glutamine, at 37°C with 5% CO_2_. For inducing the degradation of MSTO1-FLAG-mAID, 5-Ph-IAA (Cayman Chemical, 38161) was added to the culture medium at 20 μM. MSTO1 patient cells from the Cell line and DNA bank of Genetic Movement Disorders and Mitochondrial Diseases (GMD-MDbank). MSTO1 patient cells were grown in DMEM supplemented with 20% FBS (Seradigm) and Glutamax (Invitrogen).

### Cell Line Generation

The original HCT116 cell line was obtained from Jodi Nunnari, University of California Davis. To generate OsTIR1(F74G) stably expressing cell lines HCT116 cells were transfected with SpCas9-2A-GFP (#79888; Addgene) and the pAAVS1-puro (#80490; Addgene) encoding OsTIR1(F74G)-V5 and the puromycin resistance gene flanked by homology sequence targeting the AAVS1 safe harbor locus using JetPrime Polyplus (VWR Scientific). Cells were incubated in transfection reagents for 48 hrs and then grown for six days in normal media before selection in the presence of Puromycin (1 μg/mL) for 72 hours. Clonal populations were generated by plating cells at very low density, and clones were collected onto sterile filter paper dots soaked in trypsin. In clonal populations, the biallelic insertion of OsTIR1(F74G) was confirmed by genomic PCR and the expression of OsTIR1(F74G)-V5 was confirmed by Western blotting. This clonal population was used to modify the endogenous MSTO1 gene. An optimal gRNA target sequence closest to the genomic target site was chosen using CHOPCHOP and validated with Synthego CRISPR Design Tool. Individual gRNAs targeting the C-terminus of MSTO1 were cloned into SpCas9-2A-GFP. For knock-in of FLAG-AID, the donor plasmid pMK287 (#72825 Addgene) was modified to encode blasticidin resistance by replacing the hygromycin resistance cassette. gBlock designs were ordered from IDT that included a homology arm corresponding to the 3-prime end of the *MSTO1* coding sequence and a homology arm corresponding to the beginning of the 3-prime UTR of the *MSTO1* gene. The homology arms and FLAG sequences were amplified by PCR from the gBlock and cloned into the pMK287-Bsd plasmid so the homology arm composed of the 3-prime end of *MSTO1* coding sequence and FLAG was upstream of the AID sequence and downstream of AID were the FLAG and *MSTO1* 3-prime UTR homology arm sequence. These plasmids were transfected into the OsTIR1(F74G)-V5 clonal population using JetPrime PolyPlus. Cells were incubated with the transfection reagent for ∼48 hrs, grown in normal media for six days and then subject to selection with blasticidin (10 μg/mL) for ∼72 hrs. Clonal populations were generated by plating cells at very low density and clones were collected onto sterile filter paper dots soaked in trypsin. Clones were evaluated for the biallelic insertion by genomic PCR. Subsequently, the expression of the MSTO1-FLAG-mAID protein was confirmed by western blotting.

### SDS-PAGE, Western Blot analysis, and quantification

Cells were grown to ∼80% confluency, and were washed with PBS (Thermo Fisher Scientific, 20012050) before lysis with radioimmunoprecipitation assay (RIPA) buffer (25 mm Tris (pH 7.4)[Millipore Sigma], 150 mm NaCl [Fisher S2711], 1% NP-40 [Thomas Scientific 492018-50ML], 1% sodium deoxycholate [Thermo Scientific FER89904], 0.1% SDS [Fisher Scientific BP166100], 1× Halt protease inhibitor mixture [Thermo Scientific 78439]. Samples were incubated on ice for 15 min and then spun at 21,000 × *g* for 15 min at 4°C. Supernatant was transferred to a clean tube, and protein concentration was measured by BCA assay (Thermo Scientific). Following separation of 7.5 μg of whole cell lysate [or 25 μg to detect MSTO1] by SDS–PAGE, proteins were transferred to methanol activated Polyvinylidene Fluoride (PVDF) membranes. Membranes were washed in H_2_O for 10 min and then incubated in PVDF 1x TotalStain Q staining solution (#AC227; AzureBiosysetms). After decanting the staining solution, the membrane was washed in 1x TotalStain Q Wash Buffer three times for 3 minutes per wash, then visualized in the Cy3 channel. Membranes were then incubated with 4% w/v milk in tris-buffered saline (TBS) with 1% (v/v) Tween 20 (Fisher Scientific BP337-100) at room temperature for 1 to 4 hours. The indicated proteins were detected using primary rabbit or mouse antibodies incubated in 4% milk in TBST overnight at 4°C and visualized with appropriate secondary antibodies conjugated to IRDye also incubated with 4% milk in TBST for 40 minutes at room temperature (Thermo Fisher Scientific). Quantification was performed with Azure Analysis software (Azure Biosystems) and fold change calculated relative to TotalQStain (Azure Biosystems) for all figures except Figure 3D where GAPDH was used. The data were analyzed using ordinary one-way Anova with multiple comparisons or Welch’s unpaired T-test. The following antibodies were used in this study: mouse monoclonal anti-FLAG (Sigma; 1:1000); rabbit polyclonal anti-CCT1 (ThermoFisher PA5-80100; 1:1000); mouse monoclonal anti-α-tubulin (ThermoFisher Scientific clone DM1A; 1:5000); mouse monoclonal anti-β-tubulin (ThermoFisher Scientific 32-2600; 1:1000); mouse monoclonal anti-γ-tubulin (ThermoFisher Scientific MA1-850-A555; 1:1000); rabbit polyclonal anti-β-Actin (Cell Signaling 4967S; 1:1000); rabbit monoclonal anti-GAPDH (Cell Signaling (14C10); 1:500); mouse monoclonal anti-V5 (ThermoFisher R960-25; 1:1000); mouse monoclonal anti-Mfn2 (Sigma clone 4H8; 1:1000); and rabbit polyclonal anti-Mfn1 (gift from Jodi Nunnari, University of California, Davis; 1:500). Briefly, Mfn1 antiserum was raised against His6-tagged fusion proteins comprised of full-length mouse dihydrofolate reductase and an internal region of Mfn1 (residues 350–580). Fusion proteins were purified on nickel nitrilotriacetic acid columns (Thermo Fisher Scientific) in 8 M urea and eluted with 0.1% SDS and 10 mM Tris-Cl, pH 7.4.

### Microscopy

All cells were plated in no. 1.5 glass-bottomed dishes (MatTek) treated with 10 μg/mL fibronectin for 4 – 24 hrs at 37°C. HCT116 cells were incubated with 0.1 μg/ml Mitotracker Red CMX Ros, 1 μL/mL of picogreen, and 2 drops of NucBlue (Molecular Probes) for 10 min at 37°C with 5% CO_2_, then washed and incubated with complete media pre-incubated at 37°C with 5% CO_2_ for at least 4 hours before imaging. A Z-series with a step size of 0.3 μm was collected with a Nikon Ti2E confocal microscope with a 60 × NA (numerical aperture) 1.4 oil objective (Nikon), a cicero spinning disk, a 5-channel Celesta light source (Nikon), and an sCMOS camera (Prime BSI Express). Each cell line was imaged on at least three separate occasions by a blinded experimenter (n > 100 cells per experiment).

### Image analysis

Images were deconvolved using 15 iterations of 3D Landweber deconvolution. Deconvolved images were then analyzed using Nikon Elements software. Maximum intensity projections were created using Nikon Elements software and cropped in Fiji software (National Institutes of Health [NIH]). Mitochondrial morphology in mammalian cells was scored as follows: reticular indicates that fewer than 30% of the mitochondria in the cell were fragments (fragments defined as mitochondria less than or equal to 2.5 μm in length); short indicates mitochondria that were between 2.5 and 5 μm in length; fragmented indicates that most of the mitochondria in the cell were less than or equal to 2.5 μm in length. All quantifications were performed by a blinded experimenter. Data was analyzed using ordinary two-way Anova with multiple comparisons.

### Coimmunoprecipitation

Whole-cell lysates were prepared from HCT116 cells expressing MSTO1-FLAG-AID or mcherry-FLAG by incubating in lysis buffer (50 mM HEPES-KOH, pH 6.8, 300 mM NaCl, 1 mM MgCl_2_, 10% glycerol) with 0.5% NP-40 (MilliporeSigma) and 1× Halt Protease Inhibitor (Thermo Fisher Scientific) on a rocker at 4°C for 1 hour. Lysates were cleared at 21,000 × g for 30 min at 4°C and supernatant was transferred to a new tube. Supernatant was incubated with 50 μl magnetic mMACS anti-DYKDDDDK microbeads (Miltenyi Biotec) for 4 hours on a rocker at 4°C. The sample was applied to a MACS column (Miltenyi Biotec) and washed twice with lysis buffer and four times with wash buffer (50 mM HEPES-KOH, pH 6.8, 300 mM NaCl, 1 mM MgCl_2_, 10% glycerol, 0.1% NP-40 (MilliporeSigma) and 1× Halt Protease Inhibitor). One column volume (25 μl) elution buffer 50 mM HEPES-KOH, pH 6.8, 300 mM NaCl, 1 mM MgCl_2_, 10% glycerol, 0.1% NP-40 (MilliporeSigma), 6M urea, and 1× Halt Protease Inhibitor) was added and incubated for 15 min at room temperature, then proteins were eluted twice with a total of 50 μl elution buffer. Samples were run on an SDS–PAGE gel and transferred onto PVDF membranes. Membranes were blocked with 4% milk in TBST for at least 45 min and were incubated with the indicated primary antibody at 4°C overnight - anti-Mfn2, anti-Mfn1, anti-FLAG, anti-CCT1, anti-A-tubulin, anti-B-Actin, anti-CCT2 (Abcam, ab92746, 1:1000), anti-CCT3 (Santa Cruz Biotechnology, sc-271336, 1:100), anti-CCT7 (Santa Cruz Biotechnology, sc-271951, 1:143), and anti-GAPDH (Cell Signaling (14C10); 1:500). The next day membranes were incubated with DyLight secondary antibody (Invitrogen) for ∼ 40 min at room temperature; anti-CCT3 and anti-CCT7 membranes were incubated with horseradish peroxidase linked secondary antibody (Cell Signaling Technology) for 40 min at room temperature. Membranes were imaged on Azure Biosystems Sapphire Imager (Azure Biosystems).

### Genomic DNA qPCR

Total genomic DNA (gDNA) (nuclear and mitochondrial DNA) was extracted from unedited HCT116 and MSTO1-FLAG-AID cells treated with either DMSO or 20 μM 5-Ph-IAA using the DNeasy Blood and Tissue Kits for DNA Isolation (Qiagen, 69504) according to manufacturer’s instructions. Relative mtDNA copy number was analyzed by real-time quantitative PCR (qPCR) using the QuantStudio 6 Flex Real-Time PCR system (Thermo Fisher Scientific). Primer sequences specific to mtDNA and the nuclear-encoded housekeeping gene GAPDH were used. *Cytb:* 5’ TCTCCGATCCGTCCCTAACA 3’ and 5’ TGATTGGCTTAGTGGGCGAA 3’. Homo sapiens Mitochondrial gene: 5’ ACA CCC TCC TAG CCT TAC TAC 3’ and 5’ GAT ATA GGG TCG AAG CCG C 3’. *MT-ND1*: 5’ TACGGGCTACTACAACCCTTC 3’ and 5’ ATGGTAGATGTGGCGGGTTT 3’. *GAPDH*: 5’ GGCATTGCCCTCAACGACC 3’and 5’ AAAGAGTTGTCAGGGCCCTTTTTC 3’.

qPCR reactions were prepared to a total of 5 μL per reaction containing 8 μL PowerUp SYBR Green Master Mix (Thermo Fisher Scientific, A25742), 100 ng gDNA and 250 nM each forward and reverse primers. MtDNA copy number relative to GAPDH was analyzed using the delta delta Ct (ΔΔCt) method and is represented as fold change of control. Relative mtDNA copy number is presented as mean ± SD from at least three independent biological replicates and ordinary two-way ANOVA with multiple comparisons were used to determine statistical significance.

### shCCT1

shRNA target sequences were obtained using the RNAi Consortium (TRC; Broad Institute). The sequences of CCT1 short hairpin RNA (shRNA) were as follows: CCT1-shRNA/F: 5’ CCGGGGTGTACAGGTGGTCATTATTCAAGAGATAATGACCACCTGTACACCTTTTTT G 3’; and CCT1-shRNA/R: 5’ AATTCAAAAAAGGTGTACAGGTGGTCATTATCTCTTGAATAATGACCACCTGTACACC 3’. Each CCT1 shRNA and the RFP shRNA control were inserted into the pKLO.1 lentiviral vector. HEK293T cells (Cell Biolabs) were maintained in DMEM complete media supplemented and plated at ∼80% confluency the day before transfection. HEK293T cells were transfected with pKLO.1 lentiviral vector expressing CCT shRNA or RFP shRNA using FuGENE HD (Promega). Viral supernatants were collected 48 hrs post transfection and frozen in -80 C overnight. Viral supernatants were incubated with HCT116 + MSTO1-FLAG-AID cells in the presence of 8 mg/mL polybrene. Approximately 48 hrs after viral transduction, HCT116 + MSTO1-FLAG-AID cells were split, and 1 μg/ml of neomycin selection was added for 4-5 days. Cells were either seeded onto no. 1.5 glass-bottomed dishes (MatTek) coated with 10 μg/ml fibronectin (Sigma-Aldrich) for imaging 3 days later or one well of a six-well dish for whole cell lysate preparation 3 days later (described above).

### BN-PAGE

Either untreated HCT116-WT or MSTO1-AID-FLAG cells treated with DMSO or auxin were lysed in 0.5% NP-40 (MilliporeSigma), 25 mM HEPES pH 7.5, 150 mM KCl, 10% glycerol, 1.5 mM MgCl_2_, 1 mM DTT, 1× Halt Protease Inhibitor (Thermo Fisher Scientific) for 10 min in 4°C. Lysates were centrifuged at 19,0000 × *g* at 4°C for 10 min. The cleared lysate was mixed with Invitrogen NativePAGE 5% G-250 sample additive to a final concentration of 0.25%. Samples were separated on a Novex NativePAGE 4–16% Bis-Tris Protein Gels (Invitrogen) at 4°C. Gels were run at 40 V for 30 min and then 100 V for 30 min with dark cathode buffer (1× NativePAGE Running Buffer [Invitrogen], 0.02% [wt/vol] Coomassie G-250). Dark cathode buffer was replaced with light cathode buffer (1× NativePAGE Running Buffer [Invitrogen], 0.002% [wt/vol] Coomassie G-250) and the gel was run at 250 V for 60–75 min until the dye front ran off the gel. After electrophoresis was complete, gels were transferred to polyvinylidene fluoride membrane (PVDF) (Bio-Rad Laboratories) at 30 V for 16 h in transfer buffer (25 mM Tris, 192 mM glycine, 20% methanol). Membranes were subsequently incubated with 8% acetic acid for 15 min and washed with dH_2_O for 5 min. Membranes were dried at 25°C for 60 min and then rehydrated in 100% methanol and washed in dH_2_O. Membranes were blocked with 4% milk in TBST for 20 min at room temperature and were incubated with anti-FLAG, anti-CCT1, anti-CCT2, anti-CCT3, and anti-CCT7 in 4% milk/TBST for 4 hr at room temperature or overnight at 4°C. Membranes were incubated with horseradish peroxidase linked secondary antibody in 4% milk/TBST (Cell Signaling Technology) at room temperature for 1 hr. Membranes were developed in SuperSignal Femto ECL reagent (Thermo Fisher Scientific) for 5 min and imaged on Azure Biosystems Sapphire Imager (Azure Biosystems). NativeMark Unstained Protein Standard (Life Technologies) was used to estimate molecular weights of TRiC and MSTO1 protein complexes.

## Author Contributions

AMB: conceptualization, methodology, investigation, formal analysis, writing (draft and editing), visualization. CH: investigation, validation. SH: conceptualization, methodology, investigation, formal analysis, writing (draft and editing), visualization, supervision, project administration, funding acquisition.

## Acknowledgements

We thank the members of the Hoppins lab for scientific support, discussion, and advice. Funding was provided by grants from the National Institute of General Medical Sciences (R01GM118509 to SH; T32GM136534 to AMB;) and the Muscular Dystrophy Association (MDA 1053188 to SH).

---

**Figure S1. Confirmation of the genomic integration events.** (**A**) Schematic of four PCR primer pairs (P1-P4) used to confirm the integration of OsTiR1-V5 at the AAVS1 safe harbor locus. (**B**) PCR genotyping reactions from three clonal populations (MSTO1-FLAG-AID-1, -2, and -3) confirms homozygous insertion of OsTiR1-V5. (**C**) Schematic of three PCR primer pairs (P5-P7) used to confirm the integration of the AID-FLAG cassette at C terminus of *MSTO1* (P5, P6-1, P7-1) and *MSTO2P* (P5, P6-2, P7-2). Plus symbols (+) denote the location of two *MSTO2P*-specific insertions. (**D**) PCR genotyping reactions from three clonal populations (MSTO1-FLAG-AID-1, -2, and -3) confirm homozygous insertion FLAG-AID at C terminus of *MSTO1*. **(E)** PCR genotyping reactions from three clonal populations (MSTO1-FLAG-AID-1, -2, and -3) confirm homozygous insertion FLAG-AID at C terminus of *MSTO2P*. The MSTO2P-specific insertions in exon 13 result in a larger PCR product for P5. The P7-2 primers amplify both MSTO1 (green arrow) and MSTO2P (pink arrow).

**Figure S2 MSTO1 is depleted after 2 hours of auxin treatment in two additional clonal populations. (A-B)** Representative Western blot showing depletion of MSTO1-FLAG-AID. Whole cell lysate from wild-type HCT116 (WT) or the second (MSTO1-FLAG-AID-2)(A) and third (MSTO1-FLAG-AID-3)(B) clonal populations were treated with either vehicle (DMSO) or 20 *μ*M auxin or the indicated time were subject to SDS-PAGE and immunoblotting with anti-FLAG and anti-GADPH with Total StainQ used as loading control. Molecular weight markers are indicated in kDa on the left. **(C-D)** Quantification of Western blot as shown in (A and B). Fold change of MSTO1-FLAG-AID protein level relative to untreated MSTO1-FLAG-AID cells was calculated for the indicated clonal population. The graph shows the mean and standard deviation from three independent experiments, ns: not significant, ***P < 0.001, ordinary one-way Anova with multiple comparisons.

**Figure S3. MSTO1-depleted cells have fragmented mitochondria in two additional clonal populations. (A)** Representative live cell images of either MSTO1-FLAG-AID-2 or MSTO1-FLAG-AID-3 treated with DMSO (vehicle) or 20 *μ*M auxin for 6 days. Mitochondria were labeled with Mitotracker Red CMXRos, mtDNA were labeled with Quant-iT™ PicoGreen® dsDNA Reagent, and nuclei were labeled with NucBlue. All were visualized by fluorescence microscopy. Scale bar is 5 *μ*M **(B)** Quantification of mitochondrial morphology in cell lines described in (A). The graph shows mean of at least 100 cells and standard deviation from three independent experiments. ns: not significant, **P < 0.001, ordinary two-way Anova with multiple comparisons. **(C)** Quantification of mtDNA in MSTO1-AID-FLAG-2 cells treated with vehicle or 20 *μ*M auxin for 6 days. Cellular DNA samples were prepared from the indicated cell line, and mtDNA was quantified by qPCR relative to *GAPDH*, a nuclear housekeeping gene. The graph shows the mean and standard deviation of three independent experiments.

**Figure S4. CCT1 levels are significantly reduced in two additional clonal populations. (A-B)** Representative Western blot showing protein levels of CCT1 following depletion of MSTO1-FLAG-AID in the second (A) and third (B) clonal population. Whole cell lysates from wild-type HCT116 (WT) or MSTO1-FLAG-AID cells treated with either vehicle (DMSO) or 20 *μ*M auxin for the indicated time were subject to SDS-PAGE and immunoblotting with anti-CCT1, and anti-GADPH with Total StainQ used as loading control. Molecular weight markers are indicated in kDa on the left. **(C-D)** Quantification of Western blot as shown in (A and B). Fold change of CCT1 protein level relative to WT cells was calculated. The graph shows the mean and standard deviation from 3 independent experiments for the second (C) and third (D) clonal populations, **P < 0.01, ***P < 0.001, ordinary one-way Anova with multiple comparisons. **(E)** Representative Western blot showing protein levels of CCT1 at time points when changes in mitochondrial morphology were observed. Whole cell lysates from wild-type HCT116 (WT) or MSTO1-FLAG-AID-2 cells treated with either vehicle (DMSO) or 20 *μ*M auxin for the indicated time were subject to SDS–PAGE and immunoblotting with anti-CCT1, and anti-GADPH with Total StainQ used as loading control. Molecular weight markers are indicated in kDa on the left. **(F)** Quantification of Western blot as shown in (E). Fold change of CCT1 protein levels relative to WT HCT116 cells was calculated. The graph shows the mean and standard deviation from 3 independent experiments.

**Figure S5 Protein level of cytoskeletal components is not changed in MSTO1-depleted cells.** (**A**) Representative Western blot showing protein levels of cytoskeleton components following depletion of MSTO1-FLAG-AID in the second clonal population. Whole cell lysates from MSTO1-FLAG-AID-2 cells either untreated of treated with either vehicle (DMSO) or 20 *μ*M auxin for the indicated time were subject to SDS–PAGE and immunoblotting with anti-*α*-tubulin, anti-*β*-tubulin, anti-*γ*-tubulin, anti-*β*-actin, and anti-GADPH with Total StainQ as loading control. (**B**) Quantification of Western blot as shown in (A). Fold change of *α*-tubulin, *β*-tubulin, *γ*-tubulin, or *β*-actin protein levels relative to untreated cells was calculated. The graph shows the mean and standard deviation from 3 independent experiments. **(C)** Representative live cell images of MSTO1-FLAG-AID-1 cells treated with either vehicle (DMSO) or 20 *μ*M auxin 6 days. Microtubules were immunostained with anti-*α*-tubulin and actin was immunostained with phalloidin. Both were visualized by fluorescence microscopy. Scale bar is 5 *μ*M.

**Figure S6. MSTO1 interacts with TRiC in the second clonal population of MSTO1-FLAG-AID. (A)** Representative Western blot of immunoprecipitation of either control (mCherry-FLAG) or MSTO1-FLAG-AID-2 using anti-FLAG magnetic beads. Lysate fraction (Total, 0.66%) and immunoprecipitates (Elution, 6.6%) were subject to SDS-PAGE and immunoblotting with anti-FLAG, anti-Mfn1, anti-Mfn2, anti-CCT1, anti-CCT3, anti-CCT7, anti-CCT2, or anti-GADPH. **(B)** Quantification of co-immunoprecipitated CCT1, CCT3, CCT7, and CCT2 protein levels compared to total protein levels from Western blot in (A).

**Figure S7. Biological replicates of BN-PAGE experiments. (A-C)** Representative BN-PAGE of biological replicates showing fully assembled TRiC and early assembly scaffolds following depletion of MSTO1 in two clonal populations. Whole cell lysates from MSTO1-FLAG-AID-2 or MSTO1-FLAG-AID-3 cells either untreated or treated with vehicle (DMSO) or 20 *μ*M auxin for 3 days were subject to BN-PAGE followed by immunoblotting with anti-CCT2 (A), anti-CCT7 (B), or anti-CCT3 (C). Open arrowhead indicates early assembly scaffolds and closed arrowhead indicates fully assemble TRiC. Molecular weight markers are indicated in kDa on the left.

## References cited

1. S. Srivastava, The Mitochondrial Basis of Aging and Age-Related Disorders. Genes 8, 398 (2017).

2. M. Forte, et al., The role of mitochondrial dynamics in cardiovascular diseases. Br. J. Pharmacol. 178, 2060–2076 (2021).

3. M. L. Boland, A. H. Chourasia, K. F. Macleod, Mitochondrial dysfunction in cancer. Front. Oncol. 3, 292 (2013).

4. A. Gal, et al., MSTO1 is a cytoplasmic pro-mitochondrial fusion protein, whose mutation induces myopathy and ataxia in humans. EMBO Mol. Med. 9, 967–984 (2017).

5. A. Nasca, et al., A novel homozygous MSTO1 mutation in Ashkenazi Jewish siblings with ataxia and myopathy. J. Hum. Genet. (2021). 10.1038/s10038-020-00897-4.

6. A. Nasca, et al., Recessive mutations in *MSTO1* cause mitochondrial dynamics impairment, leading to myopathy and ataxia. Hum. Mutat. 38, 970–977 (2017).

7. D. Ardicli, et al., A novel case of MSTO1 gene related congenital muscular dystrophy with progressive neurological involvement. Neuromuscul. Disord. NMD 29, 448–455 (2019).

8. K. Iwama, et al., Novel recessive mutations in MSTO1 cause cerebellar atrophy with pigmentary retinopathy. J. Hum. Genet. 63, 263–270 (2018).

9. S. Donkervoort, et al., MSTO1 mutations cause mtDNA depletion, manifesting as muscular dystrophy with cerebellar involvement. Acta Neuropathol. (Berl.) 138, 1013–1031 (2019).

10. L. Schultz-Rogers, et al., Novel biallelic variants in MSTO1 associated with mitochondrial myopathy. Cold Spring Harb. Mol. Case Stud. 5, a004309 (2019).

11. K. Li, R. Jin, X. Wu, Whole-exome sequencing identifies rare compound heterozygous mutations in the MSTO1 gene associated with cerebellar ataxia and myopathy. Eur. J. Med. Genet. 63, 103623 (2020).

12. L. Liu, et al., Case Report: Evidences of myasthenia and cerebellar atrophy in a chinese patient with novel compound heterozygous MSTO1 variants. Front. Genet. 13, 947886 (2022).

13. B. Mao, et al., Congenital muscular dystrophies and myopathies: the leading cause of genetic muscular disorders in eleven Chinese families. BMC Musculoskelet. Disord. 26, 51 (2025).

14. R. Sharma, et al., MSTO1-related mitochondrial myopathy and ataxia syndrome: Case series and literature review. Neuromuscul. Disord. NMD 60, 106364 (2026).

15. V. Mottier-Pavie, G. Cenci, F. Vernì, M. Gatti, S. Bonaccorsi, Phenotypic analysis of misato function reveals roles of noncentrosomal microtubules in Drosophila spindle formation. J. Cell Sci. 124, 706–717 (2011).

16. V. Palumbo, et al., Misato Controls Mitotic Microtubule Generation by Stabilizing the TCP-1 Tubulin Chaperone Complex. Curr. Biol. 25, 1777–1783 (2015).

17. S. Min, W. Yoon, H. Cho, J. Chung, Misato underlies visceral myopathy in Drosophila. Sci. Rep. 7, 17700 (2017).

18. W. Ji, A. L. Hatch, R. A. Merrill, S. Strack, H. N. Higgs, Actin filaments target the oligomeric maturation of the dynamin GTPase Drp1 to mitochondrial fission sites. eLife 4, e11553 (2015).

19. R. Chakrabarti, et al., INF2-mediated actin polymerization at the ER stimulates mitochondrial calcium uptake, inner membrane constriction, and division. J. Cell Biol. 217, 251–268 (2018).

20. A. S. Moore, E. L. F. Holzbaur, Mitochondrial-cytoskeletal interactions: dynamic associations that facilitate network function and remodeling. Curr. Opin. Physiol. 3, 94–100 (2018).

21. D. Gestaut, A. Limatola, L. Joachimiak, J. Frydman, The ATP-powered gymnastics of TRiC/CCT: an asymmetric protein folding machine with a symmetric origin story. Curr. Opin. Struct. Biol. 55, 50–58 (2019).

22. D. Gestaut, et al., Structural visualization of the tubulin folding pathway directed by human chaperonin TRiC/CCT. Cell 185, 4770–4787.e20 (2022).

23. P. Gatti, C. Schiavon, J. Cicero, U. Manor, M. Germain, Mitochondria- and ER-associated actin are required for mitochondrial fusion. Nat. Commun. 16, 451 (2025).

24. A. Y. Dunn, M. W. Melville, J. Frydman, Review: cellular substrates of the eukaryotic chaperonin TRiC/CCT. J. Struct. Biol. 135, 176–184 (2001).

25. A. Camasses, A. Bogdanova, A. Shevchenko, W. Zachariae, The CCT chaperonin promotes activation of the anaphase-promoting complex through the generation of functional Cdc20. Mol. Cell 12, 87–100 (2003).

26. J. R. Zebol, et al., The CCT/TRiC chaperonin is required for maturation of sphingosine kinase 1. Int. J. Biochem. Cell Biol. 41, 822–827 (2009).

27. A. Freund, et al., Proteostatic control of telomerase function through TRiC-mediated folding of TCAB1. Cell 159, 1389–1403 (2014).

28. D. W. Neef, et al., A direct regulatory interaction between chaperonin TRiC and stress-responsive transcription factor HSF1. Cell Rep. 9, 955–966 (2014).

29. S. V. Antonova, et al., Chaperonin CCT checkpoint function in basal transcription factor TFIID assembly. Nat. Struct. Mol. Biol. 25, 1119–1127 (2018).

30. J. Cuéllar, et al., Structural and functional analysis of the role of the chaperonin CCT in mTOR complex assembly. Nat. Commun. 10, 2865 (2019).

31. Y. Cong, et al., 4.0-Å resolution cryo-EM structure of the mammalian chaperonin TRiC/CCT reveals its unique subunit arrangement. Proc. Natl. Acad. Sci. 107, 4967–4972 (2010).

32. A. Leitner, et al., The Molecular Architecture of the Eukaryotic Chaperonin TRiC/CCT. Structure 20, 814–825 (2012).

33. S. Reissmann, et al., A Gradient of ATP Affinities Generates an Asymmetric Power Stroke Driving the Chaperonin TRIC/CCT Folding Cycle. Cell Rep. 2, 866–877 (2012).

34. J. M. Archibald, C. Blouin, W. F. Doolittle, Gene duplication and the evolution of group II chaperonins: implications for structure and function. J. Struct. Biol. 135, 157–169 (2001).

35. K. Betancourt Moreira, et al., A hierarchical assembly pathway directs the unique subunit arrangement of TRiC/CCT. Mol. Cell 83, 3123–3139.e8 (2023).

36. C. Zeng, et al., Revisiting the chaperonin T-complex protein-1 ring complex in human health and disease: A proteostasis modulator and beyond. Clin. Transl. Med. 14, e1592 (2024).

37. Y. Yagita, E. Zavodszky, S.-Y. Peak-Chew, R. S. Hegde, Mechanism of orphan subunit recognition during assembly quality control. Cell 186, 3443–3459.e24 (2023).

38. O. A. Sergeeva, C. Haase-Pettingell, J. A. King, Co-expression of CCT subunits hints at TRiC assembly. Cell Stress Chaperones 24, 1055–1065 (2019).

39. M. P. Collier, et al., Native mass spectrometry analyses of chaperonin complex TRiC/CCT reveal subunit N-terminal processing and re-association patterns. Sci. Rep. 11, 13084 (2021).

40. O. A. Sergeeva, et al., Human CCT4 and CCT5 chaperonin subunits expressed in Escherichia coli form biologically active homo-oligomers. J. Biol. Chem. 288, 17734–17744 (2013).

41. A. R. Kusmierczyk, M. Hochstrasser, Some assembly required: dedicated chaperones in eukaryotic proteasome biogenesis. Biol. Chem. 389, 1143–1151 (2008).

42. K. Kato, T. Satoh, Structural insights on the dynamics of proteasome formation. Biophys. Rev. 10, 597–604 (2018).

43. A. Rousseau, A. Bertolotti, Regulation of proteasome assembly and activity in health and disease. Nat. Rev. Mol. Cell Biol. 19, 697–712 (2018).

44. J. Baßler, E. Hurt, Eukaryotic Ribosome Assembly. Annu. Rev. Biochem. 88, 281–306 (2019).

45. K. Dörner, C. Ruggeri, I. Zemp, U. Kutay, Ribosome biogenesis factors-from names to functions. EMBO J. 42, e112699 (2023).

46. A. Vanden Broeck, S. Klinge, Eukaryotic Ribosome Assembly. Annu. Rev. Biochem. 93, 189–210 (2024).

47. A. Yesbolatova, T. Natsume, K.-I. Hayashi, M. T. Kanemaki, Generation of conditional auxin-inducible degron (AID) cells and tight control of degron-fused proteins using the degradation inhibitor auxinole. Methods San Diego Calif 164-165, 73–80 (2019).

48. A. Yesbolatova, et al., The auxin-inducible degron 2 technology provides sharp degradation control in yeast, mammalian cells, and mice. Nat. Commun. 11, 5701 (2020).

49. Y. Saito, M. T. Kanemaki, Targeted Protein Depletion Using the Auxin-Inducible Degron 2 (AID2) System. Curr. Protoc. 1, e219 (2021).

50. V. Yu. Kuryshev, et al., An anthropoid-specific segmental duplication on human chromosome 1q22. Genomics 88, 143–151 (2006).

51. M. Kimura, Y. Okano, Human Misato regulates mitochondrial distribution and morphology. Exp. Cell Res. 313, 1393–1404 (2007).

52. H. Ghozlan, A. Cox, D. Nierenberg, S. King, A. R. Khaled, The TRiCky Business of Protein Folding in Health and Disease. Front. Cell Dev. Biol. 10, 906530 (2022).

53. H. Kim, J. Park, S.-H. Roh, The structural basis of eukaryotic chaperonin TRiC/CCT: Action and folding. Mol. Cells 47, 100012 (2024).

54. I. R. Boldogh, L. A. Pon, Mitochondria on the move. Trends Cell Biol. 17, 502–510 (2007).

55. A. J. Kruppa, F. Buss, Motor proteins at the mitochondria-cytoskeleton interface. J. Cell Sci. 134, jcs226084 (2021).

56. J. L. Bocanegra, et al., The MyMOMA domain of MYO19 encodes for distinct Miro-dependent and Miro-independent mechanisms of interaction with mitochondrial membranes. Cytoskeleton 77, 149–166 (2020).

57. A. S. Moore, et al., Actin cables and comet tails organize mitochondrial networks in mitosis. Nature 591, 659–664 (2021).

58. O. Sato, et al., Mitochondria-associated myosin 19 processively transports mitochondria on actin tracks in living cells. J. Biol. Chem. 298, 101883 (2022).

59. S. M. Coscia, et al., Myo19 tethers mitochondria to endoplasmic reticulum-associated actin to promote mitochondrial fission. J. Cell Sci. 136, jcs260612 (2023).

60. S. M. Coscia, et al., An interphase actin wave promotes mitochondrial content mixing and organelle homeostasis. Nat. Commun. 15, 3793 (2024).

61. F. Korobova, V. Ramabhadran, H. N. Higgs, An actin-dependent step in mitochondrial fission mediated by the ER-associated formin INF2. Science 339, 464–467 (2013).

62. A. L. Hatch, W.-K. Ji, R. A. Merrill, S. Strack, H. N. Higgs, Actin filaments as dynamic reservoirs for Drp1 recruitment. Mol. Biol. Cell 27, 3109–3121 (2016).

63. N. H. Cho, et al., OpenCell: Endogenous tagging for the cartography of human cellular organization. Science 375, eabi6983 (2022).

64. H. Schägger, G. von Jagow, Blue native electrophoresis for isolation of membrane protein complexes in enzymatically active form. Anal. Biochem. 199, 223–231 (1991).

65. X. Liu, D. Weaver, O. Shirihai, G. Hajnóczky, Mitochondrial “kiss-and-run”: interplay between mitochondrial motility and fusion-fission dynamics. EMBO J. 28, 3074–3089 (2009).

66. K. C. Stein, A. Kriel, J. Frydman, Nascent Polypeptide Domain Topology and Elongation Rate Direct the Cotranslational Hierarchy of Hsp70 and TRiC/CCT. Mol. Cell 75, 1117–1130.e5 (2019).

67. F. Rüßmann, et al., Folding of large multidomain proteins by partial encapsulation in the chaperonin TRiC/CCT. Proc. Natl. Acad. Sci. U. S. A. 109, 21208–21215 (2012).

68. Q. Zhao, et al., TRiC folds the giant ciliary protein IFT172 via a non-canonical open-state mechanism. [Preprint] (2026). Available at: https://www.biorxiv.org/content/10.64898/2026.03.28.714460v1 [Accessed 28 April 2026].

69. Y. Y. Yamamoto, et al., Asymmetry in the function and dynamics of the cytosolic group II chaperonin CCT/TRiC. PloS One 12, e0176054 (2017).

70. M. Jin, et al., An ensemble of cryo-EM structures of TRiC reveal its conformational landscape and subunit specificity. Proc. Natl. Acad. Sci. U. S. A. 116, 19513–19522 (2019).

71. K. Machida, et al., Reconstitution of the human chaperonin CCT by co-expression of the eight distinct subunits in mammalian cells. Protein Expr. Purif. 82, 61–69 (2012).

72. M. P. Collier, et al., Native mass spectrometry analyses of chaperonin complex TRiC/CCT reveal subunit N-terminal processing and re-association patterns. Sci. Rep. 11, 13084 (2021).

73. H. M. Schnell, R. M. Walsh, S. Rawson, J. Hanna, Chaperone-mediated assembly of the proteasome core particle - recent developments and structural insights. J. Cell Sci. 135, jcs259622 (2022).

74. F. Kraft, et al., Brain malformations and seizures by impaired chaperonin function of TRiC. Science 386, 516–525 (2024).

