## Supplemental Figures for "MSTO1 functions as a TRiC assembly factor linking cytosolic proteostasis to mitochondrial function"

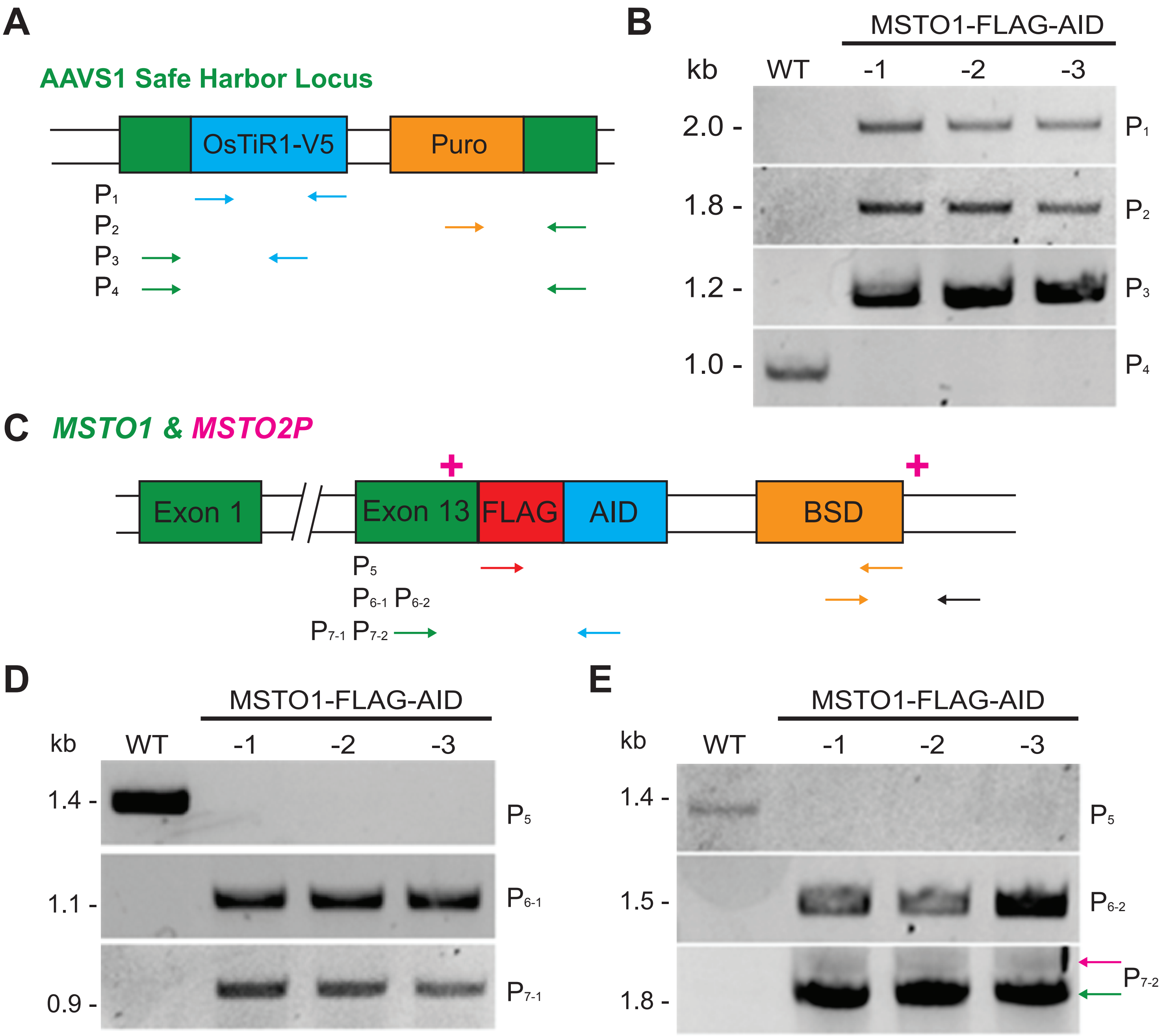


**Figure S1. Confirmation of the genomic integration events. (A)** Schematic of four PCR primer pairs (P1-P4) used to confirm the integration of OsTiR1-F74G-V5 at the AAVS1 safe harbor locus. **(B)** PCR genotyping reactions from three clonal populations (MSTO1-FLAG-AID-1, -2, and -3) confirms homozygous insertion of OsTiR1-F74G-V5. **(C)** Schematic of three PCR primer pairs (P5-P7) used to confirm the integration of the AID-FLAG cassette at C terminus of *MSTO1* (P5, P6-1, P7-1) and *MSTO2P* (P5, P6-2, P7-2). Plus symbols (+) denote the location of two *MSTO2P*-specific insertions. **(D)** PCR genotyping reactions from three clonal populations (MSTO1-FLAG-AID-1, -2, and -3) confirm homozygous insertion AID-FLAG at C terminus of *MSTO1*. **(E)** PCR genotyping reactions from three clonal populations (MSTO1-FLAG-AID-1, -2, and -3) confirm homozygous insertion AID-FLAG at C terminus of *MSTO2P*. The *MSTO2P*-specific insertion in exon 13 result in a larger product for P5 and P7-2 primers amplify both *MSTO1* (green arrow) and *MSTO2P* (pink arrow).

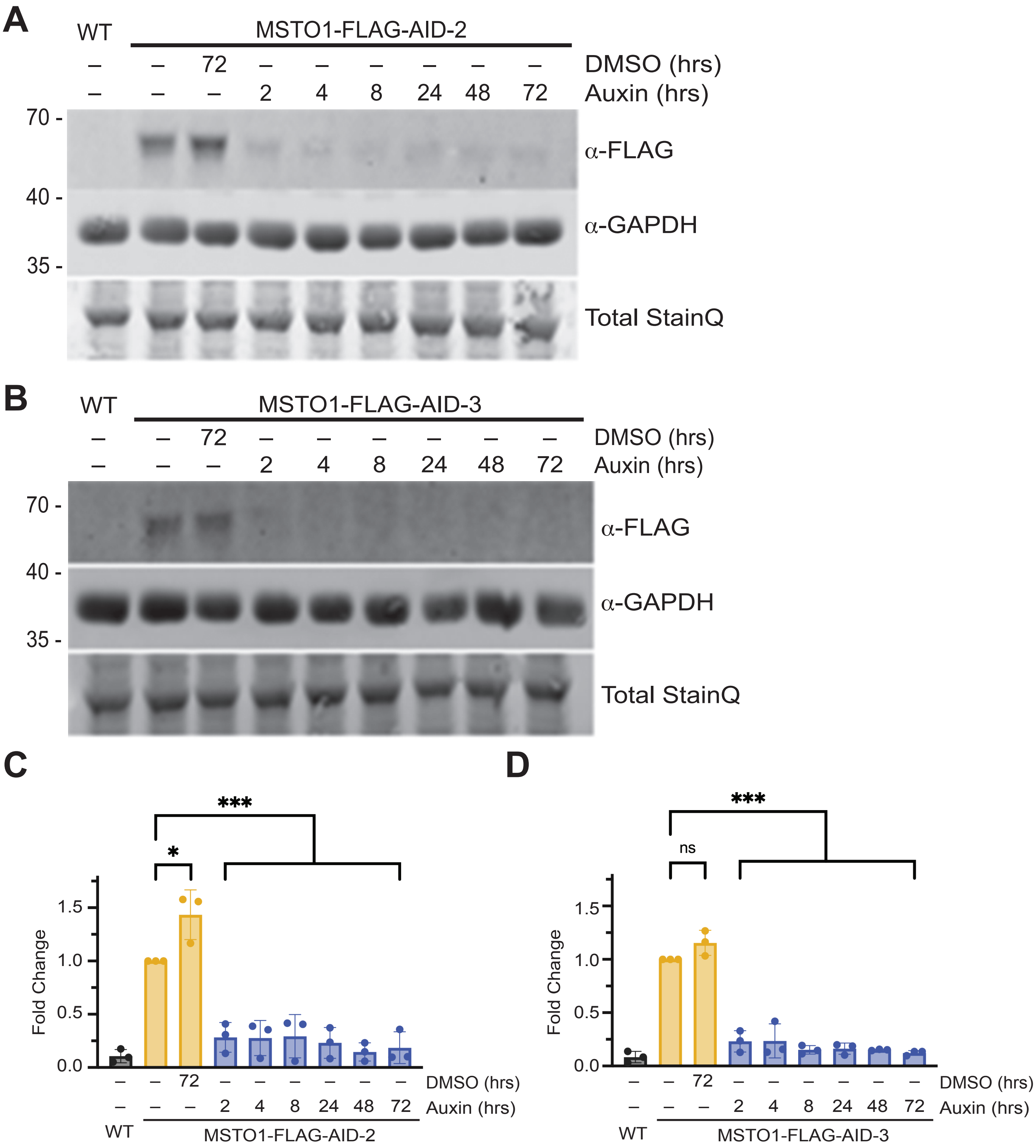


**Figure S2 MSTO1 is depleted after 2 hours of auxin treatment in two additional clonal populations. (A-B)** Representative Western blot showing depletion of MSTO1-FLAG-AID. Whole cell lysate from wild-type HCT116 (WT) or the second (MSTO1-FLAG-AID-2)(A) and third (MSTO1-FLAG-AID-3)(B) clonal populations were treated with either vehicle (DMSO) or 20 μM auxin or the indicated time were subject to SDS-PAGE and immunoblotting with anti-FLAG and anti-GADPH with Total StainQ used as loading control. Molecular weight markers are indicated in kDa on the left. **(C-D)** Quantification of Western blot as shown in (A and B). Fold change of MSTO1-FLAG-AID protein level relative to untreated MSTO1-FLAG-AID cells was calculated for the indicated clonal population. The graph shows the mean and standard deviation from three independent experiments, ns: not significant, ***$P < 0.001$, ordinary one-way Anova with multiple comparisons.

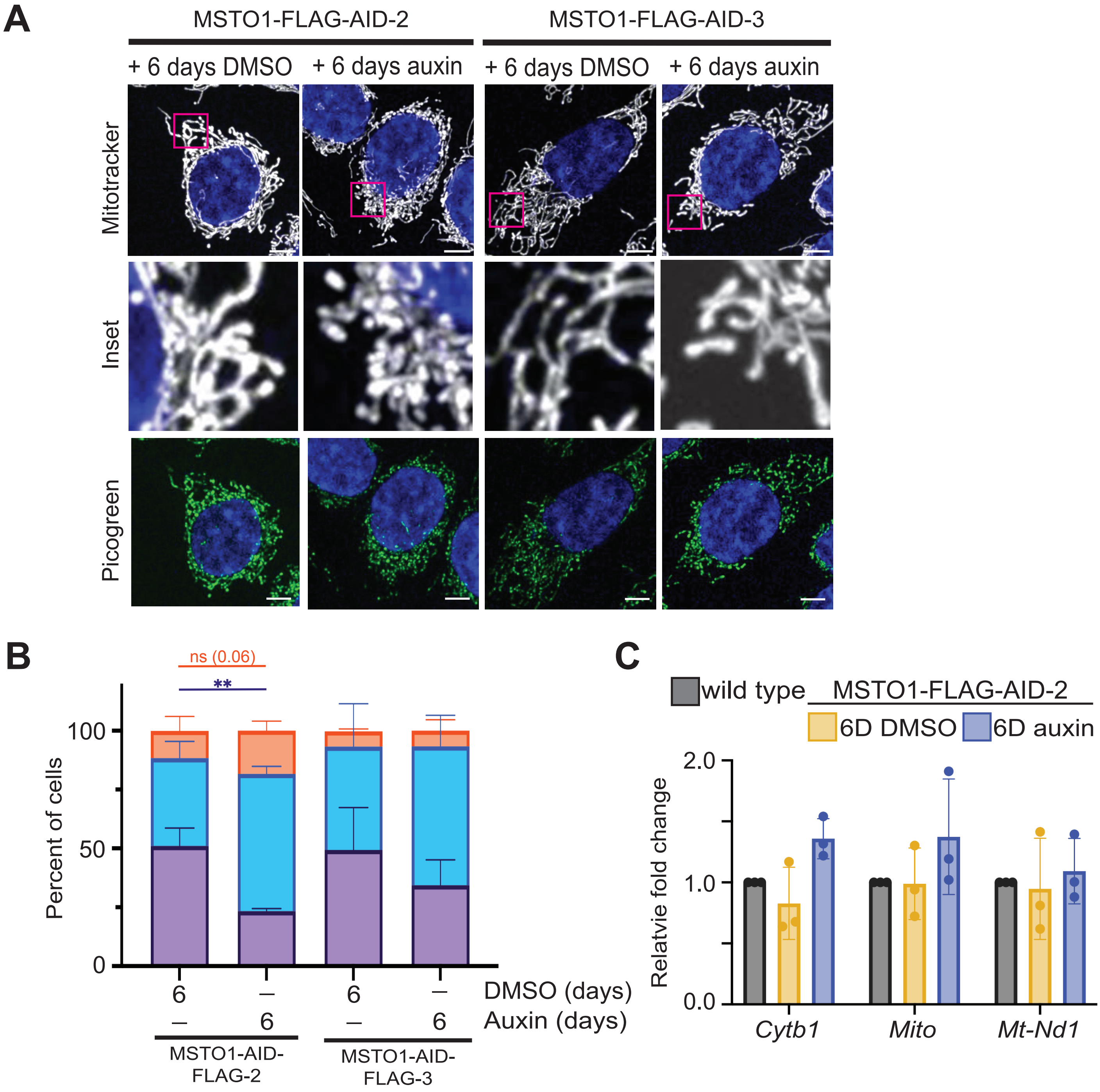


**Figure S3. MSTO1-depleted cells have fragmented mitochondria in two additional clonal populations.** (A) Representative live cell images of either MSTO1-FLAG-AID-2 or MSTO1-FLAG-AID-3 treated with DMSO (vehicle) or 20 μM auxin for 6 days. Mitochondria were labeled with Mitotracker Red CMXRos, mtDNA were labeled with Quant-iT™ PicoGreen® dsDNA Reagent, and nuclei were labeled with NucBlue. All were visualized by fluorescence microscopy. Scale bar is 5 μM. (B) Quantification of mitochondrial morphology in cell lines described in (A). The graph shows mean of at least 100 cells and standard deviation from three independent experiments. ns: not significant, $^{**}P < 0.001$, ordinary two-way Anova with multiple comparisons. (C) Quantification of mtDNA in MSTO1-AID-FLAG-2 cells treated with vehicle (DMSO) or 20 μM auxin for 6 days. Cellular DNA samples were prepared from the indicated cell line and mtDNA was quantified by qPCR relative to GAPDH, a nuclear housekeeping gene. The graph shows the mean and standard deviation of three independent experiments.

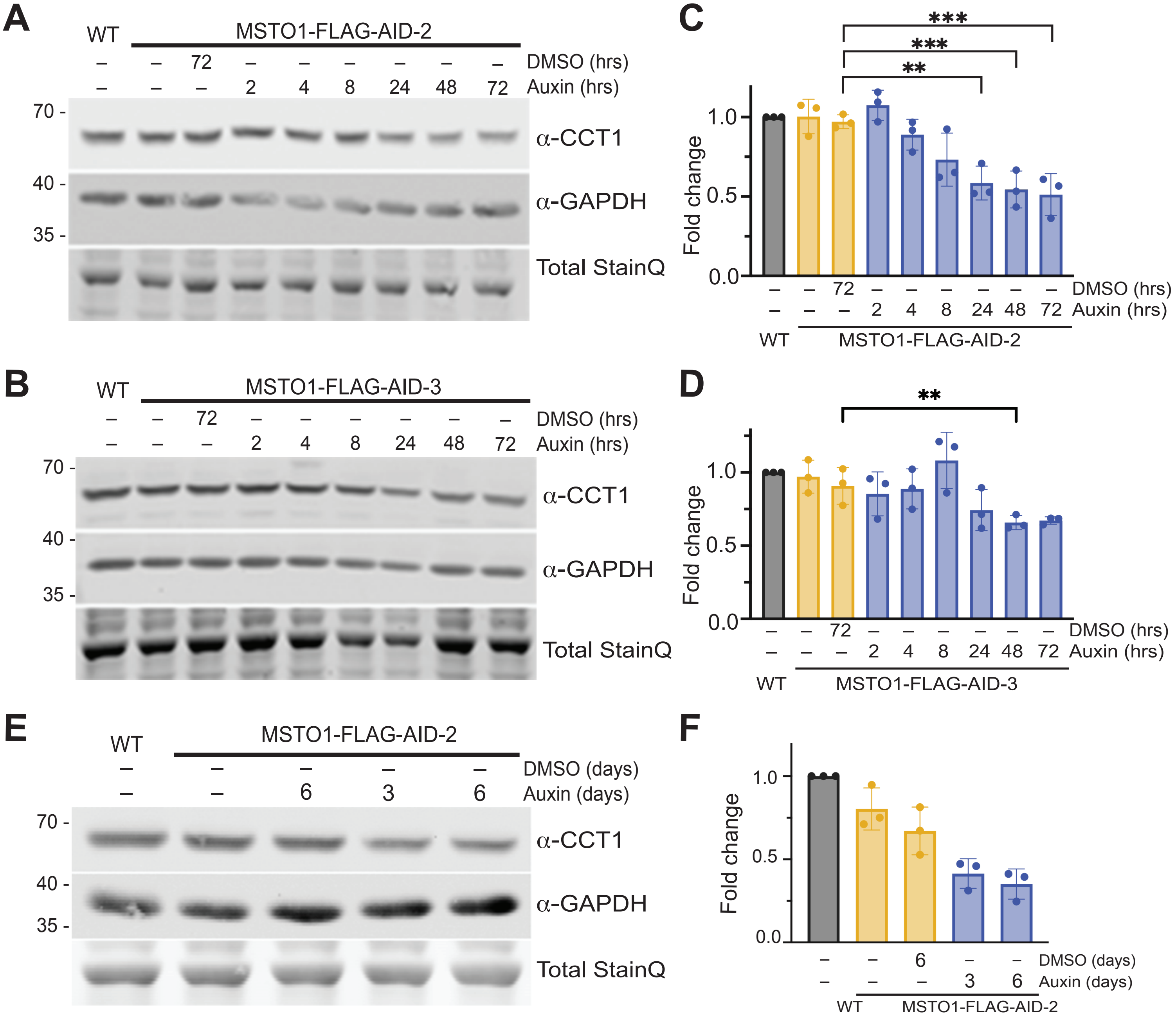


**Figure S4. CCT1 levels are significantly reduced in two additional clonal populations. (A-B)** Representative Western blot showing protein levels of CCT1 following depletion of MSTO1-FLAG-AID in the second (A) and third (B) clonal population. Whole cell lysates from wild-type HCT116 (WT) or MSTO1-FLAG-AID cells treated with either vehicle (DMSO) or 20 μM auxin for the indicated time were subject to SDS-PAGE and immunoblotting with anti-CCT1, and anti-GADPH with Total StainQ used as loading control. Molecular weight markers are indicated in kDa on the left. **(C-D)** Quantification of Western blot as shown in (A and B). Fold change of CCT1 protein level relative to WT cells was calculated. The graph shows the mean and standard deviation from 3 independent experiments for the second (C) and third (D) clonal populations, $^{**}P < 0.01$, $^{***}P < 0.001$, ordinary one-way Anova with multiple comparisons. **(E)** Representative Western blot showing protein levels of CCT1 at time points when changes in mitochondrial morphology were observed. Whole cell lysates from wild-type HCT116 (WT) or MSTO1-FLAG-AID-2 cells treated with either vehicle (DMSO) or 20 μM auxin for the indicated time were subject to SDS–PAGE and immunoblotting with anti-CCT1, and anti-GADPH with Total StainQ used as loading control. Molecular weight markers are indicated in kDa on the left. **(F)** Quantification of Western blot as shown in (E). Fold change of CCT1 protein levels relative to WT HCT116 cells was calculated. The graph shows the mean and standard deviation from 3 independent experiments.

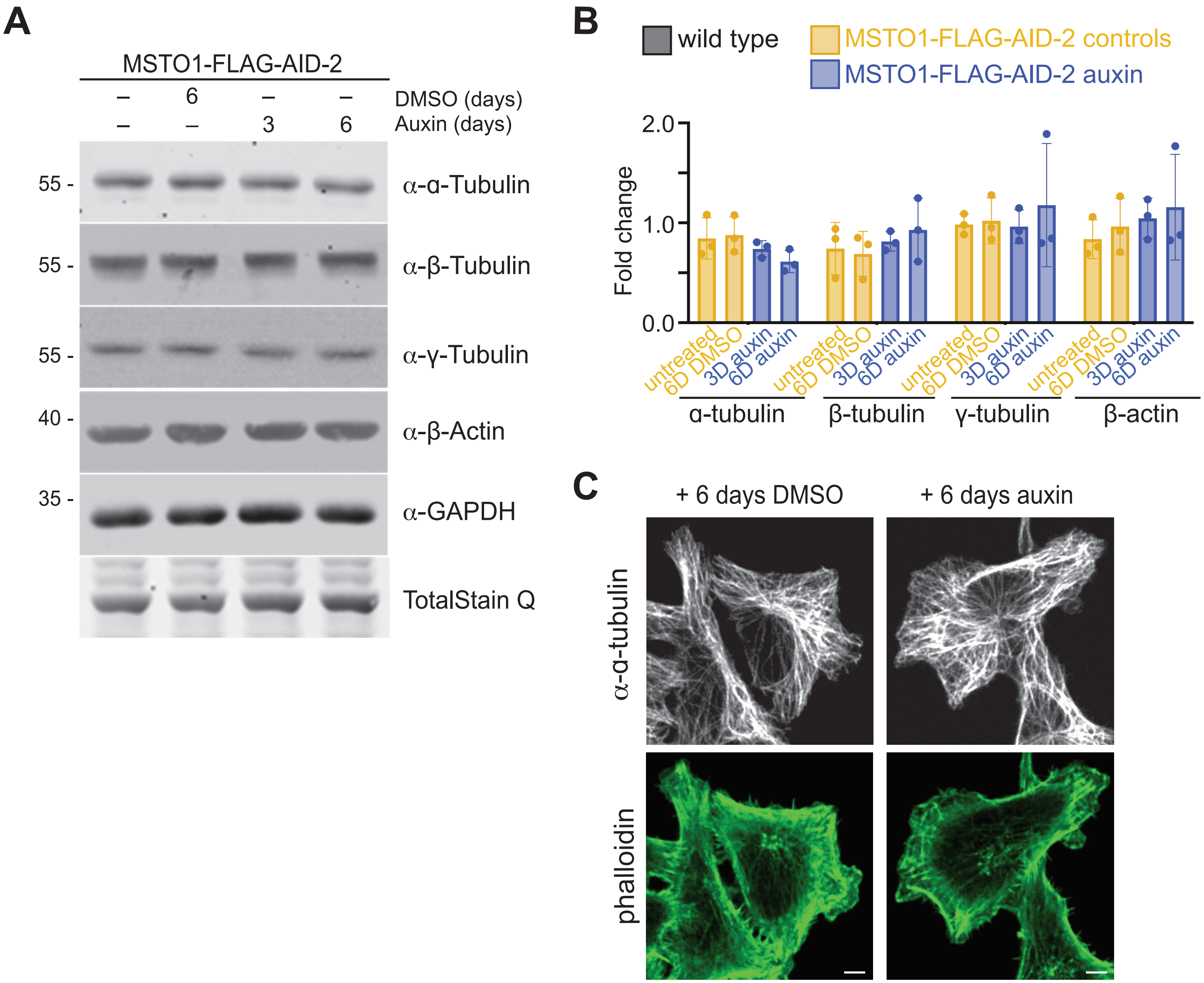


**Figure S5 Protein level of cytoskeletal components is not changed in MSTO1-depleted cells. (A)** Representative Western blot showing protein levels of cytoskeleton components following depletion of MSTO1-FLAG-AID in the second clonal population. Whole cell lysates from MSTO1-FLAG-AID-2 cells either untreated of treated with either vehicle (DMSO) or 20 μM auxin for the indicated time were subject to SDS–PAGE and immunoblotting with anti-α-tubulin, anti-β-tubulin, anti-γ-tubulin, anti-β-actin, and anti-GADPH with Total StainQ as loading control. Molecular weight markers are indicated in kDa on the left. **(B)** Quantification of Western blot as shown in (A). Fold change of α-tubulin, β-tubulin, γ-tubulin, or β-actin protein levels relative to untreated cells was calculated. The graph shows the mean and standard deviation from 3 independent experiments. **(C)** Representative live cell images of MSTO1-FLAG-AID-1 cells treated with either vehicle (DMSO) or 20 μM auxin 6 days. Microtubules were immunostained with anti-α-tubulin and actin was immunostained with phalloidin. Both were visualized by fluorescence microscopy. Scale bar is 5 μM.

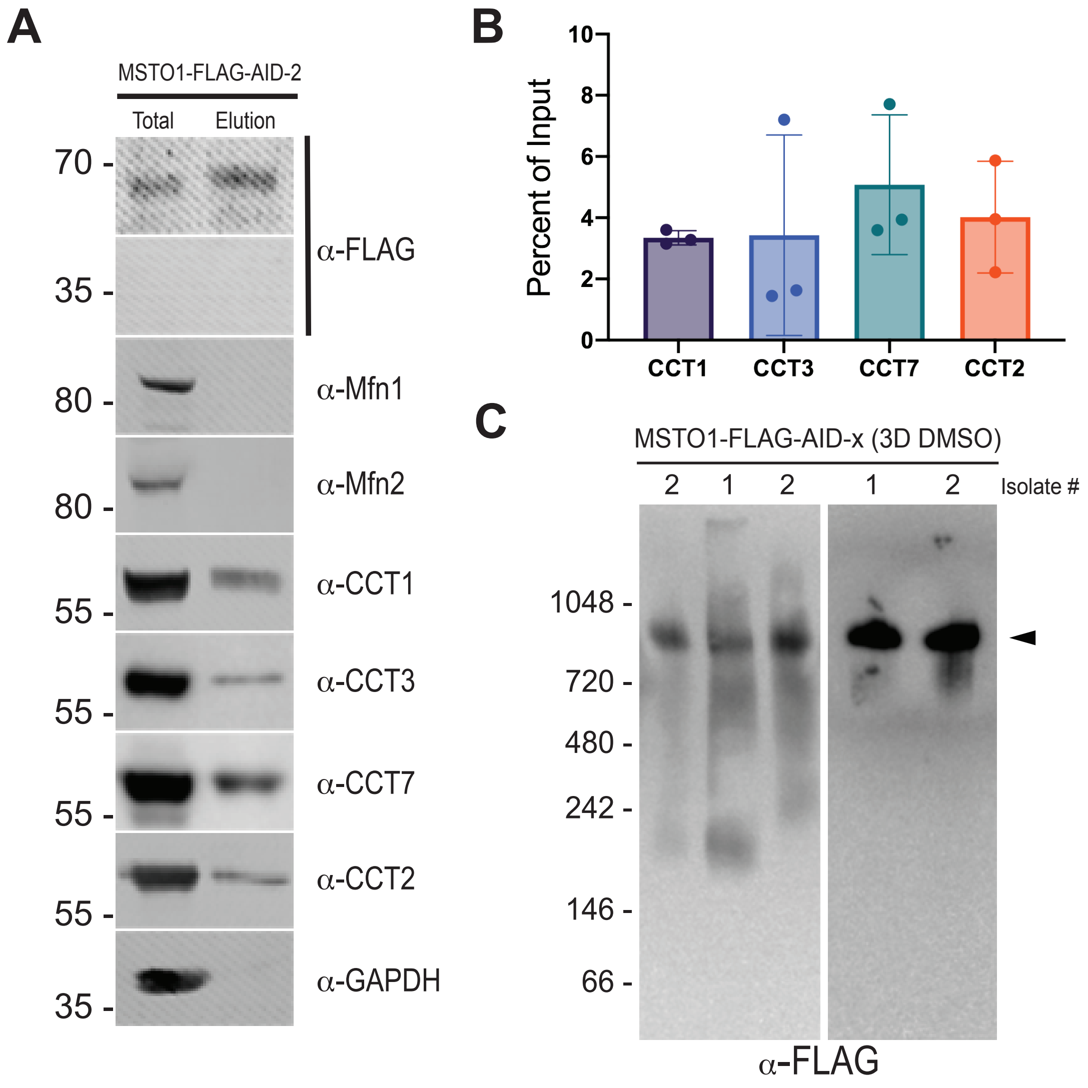


**Figure S6. Replication of MSTO1 interaction with TRiC in MSTO1-FLAG-AID-2.** **(A)** Representative Western blot of immunoprecipitation of either control (mCherry-FLAG) or MSTO1-FLAG-AID-2 using anti-FLAG magnetic beads. Lysate fraction (Total, 0.66%) and immunoprecipitates (Elution, 6.6%) were subject to SDS-PAGE and immunoblotting with anti-FLAG, anti-Mfn1, anti-Mfn2, anti-CCT1, anti-CCT3, anti-CCT7, anti-CCT2, or anti-GADPH. Molecular weight markers are indicated in kDa on the left. **(B)** Quantification of co-immunoprecipitated CCT1, CCT3, CCT7, and CCT2 protein levels compared to total protein levels from Western blot in (A). **(C)** Representative blue native polyacrylamide gel electrophoresis (BN-PAGE) of MSTO1-FLAG-AID-1 and MSTO1-FLAG-AID-2, as indicated. Whole cell lysates from MSTO1-FLAG-AID cells treated vehicle for 3 days were subject to BN-PAGE followed by immunoblotting with anti-FLAG. Molecular weight markers are indicated in kDa on the left.

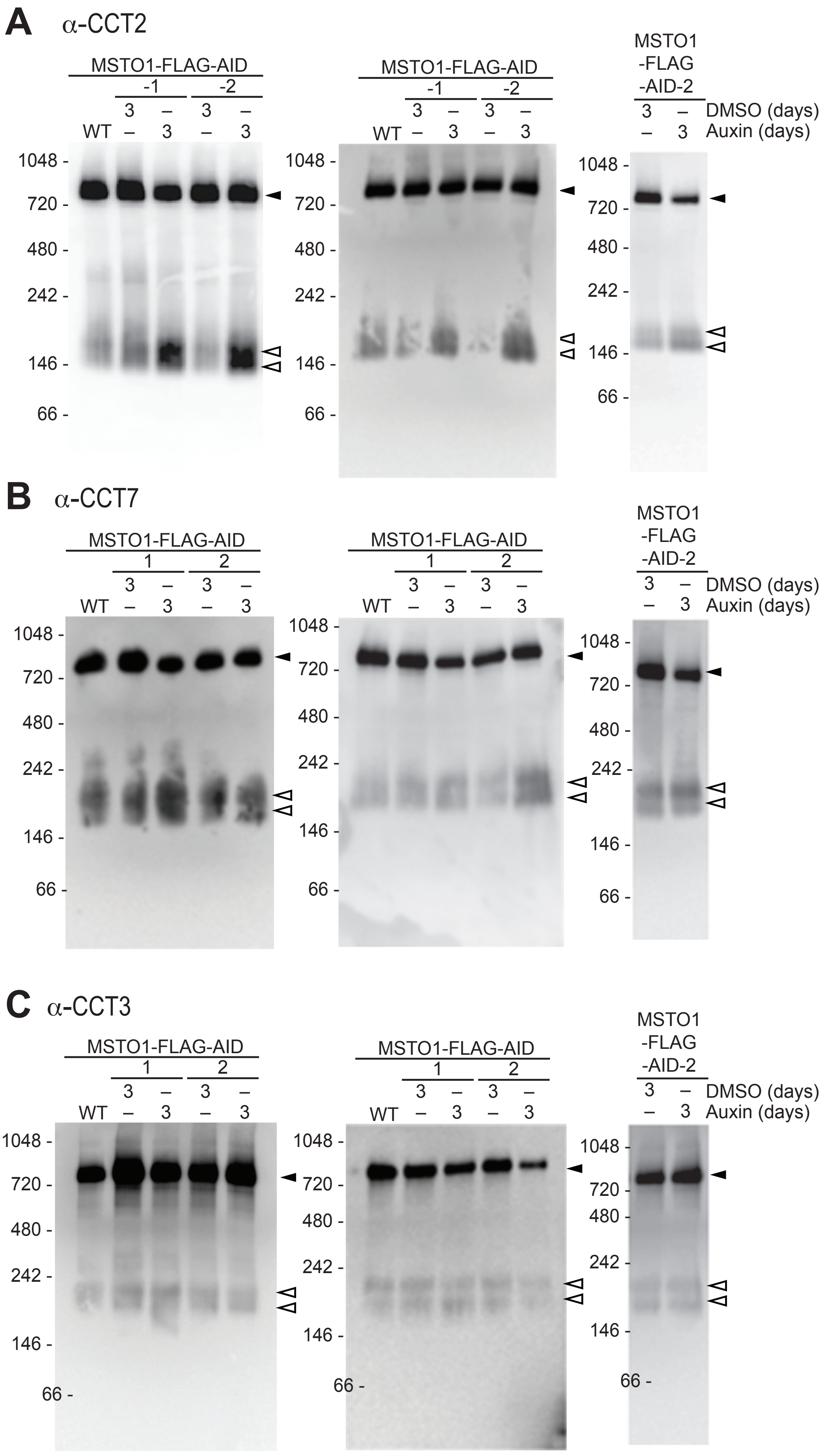


**Figure S7. Biological replicates of BN-PAGE experiments. (A-C)** Representative BN-PAGE of biological replicates showing fully assembled TRiC and early assembly scaffolds following depletion of MSTO1-FLAG-AID in two clonal populations (-1 and -2). Whole cell lysates from MSTO1-FLAG-AID cells either untreated or treated with vehicle (DMSO) or 20 μM auxin for 3 days were subject to BN-PAGE followed by immunoblotting with anti-CCT2 (A), anti-CCT3 (B), or anti-CCT7 (C). Open arrowhead indicates early assembly scaffolds and closed arrowhead indicates fully assemble TRiC. Molecular weight markers are indicated in kDa on the left.
